# Multi-laboratory Study Establishes Reproducible Methods for Plant-Microbiome Research in Fabricated Ecosystems

**DOI:** 10.1101/2024.10.02.615924

**Authors:** Vlastimil Novak, Peter F. Andeer, Eoghan King, Jacob Calabria, Connor Fitzpatrick, Jana M. Kelm, Kathrin Wippel, Suzanne M. Kosina, Chris Daum, Matt Zane, Archana Yadav, Mingfei Chen, Dor Russ, Catharine A. Adams, Trenton K. Owens, Bradie Lee, Yezhang Ding, Zineb Sordo, Romy Chakraborty, Simon Roux, Adam M. Deutschbauer, Daniela Ushizima, Karsten Zengler, Borjana Arsova, Jeffery L. Dangl, Paul Schulze-Lefert, Michelle Watt, John P. Vogel, Trent R. Northen

## Abstract

Inter-laboratory replicability is crucial yet challenging in microbiome research. Leveraging microbiomes to promote soil health and plant growth requires understanding underlying molecular mechanisms using reproducible experimental systems. In a global collaborative effort involving five laboratories, we aimed to help advance reproducibility in microbiome studies by testing our ability to replicate synthetic community assembly experiments. Our study compared fabricated ecosystems constructed using two different synthetic bacterial communities, the model grass *Brachypodium distachyon*, and sterile EcoFAB 2.0 devices. All participating laboratories observed consistent inoculum-dependent changes in plant phenotype, root exudate composition, and final bacterial community structure where *Paraburkholderia* sp. OAS925 could dramatically shift microbiome composition. Comparative genomics and exudate utilization linked the pH-dependent colonization ability of *Paraburkholderia*, which was further confirmed with motility assays. The study provides detailed protocols, benchmarking datasets, and best practices to help advance replicable science and inform future multi-laboratory reproducibility studies.

## 1. Introduction

As recent perspective papers have highlighted, establishing model microbiomes is a pressing need in environmental microbiology ^1,2^. Several years ago, a vision was presented for developing and validating standardized ‘fabricated ecosystems’ to enable replicable studies of microbiomes in ecologically relevant contexts, akin to the adoption of shared model organisms ^1^. A fabricated ecosystem is defined as a closed laboratory ecological system where all biotic and abiotic factors are initially specified/controlled. Synthetic microbial communities (SynComs) are valuable tools for bridging the gap between natural communities and studies involving axenic cultures and isolates ^3^. By limiting complexity yet retaining functional diversity and microbe-microbe interactions, SynComs can be used to unravel mechanisms underlying complex interactions, providing critical insights into community assembly processes, microbial interactions, and host physiology, e.g., plant host ^3–6^. These interactions between the host and its microbes define the holobiont concept, where the plant and its microbiome form a single dynamic ecological unit ^7^. However, standardization is essential to fully leverage the potential of SynComs and achieve replicable plant microbiome studies ^8^. This requires overcoming several challenges, including the availability of strains and standardized protocols for their growth in the laboratory. To address these challenges, we recently developed a standardized model community of 17 bacterial isolates from grass rhizosphere available through a public biobank (DSMZ), along with cryopreservation and resuscitation protocols ^9^.

Other aspects to enable replicable microbiome studies must be standardized, including sterile habitats and protocols for sample collection and analysis ^1^. As initial steps towards this vision, we developed a first-generation sterile container for fabricated ecosystems (EcoFAB device) and performed a multi-laboratory study demonstrating the reproducible physiology of the model grass *Brachypodium distachyon* ^10^. Recently, it was found that *Paraburkholderia* sp. OAS925 dominated other members of the model 17-member SynCom for *Brachypodium distachyon* root colonization^11^. Additionally, we have since developed an improved (EcoFAB 2.0 device that enables highly reproducible plant growth ^12^. The next step towards standardization is to test the replicability of microbiome formation, plant responses to microbiomes, and root exudation using these standardized laboratory habitats and SynComs. This can be achieved through inter-laboratory comparison studies or ring trials—a powerful tool in proficiency testing of analytical methods ^13,14^ that are currently underutilized in microbiome research.

Here, we describe a five-laboratory international ring trial investigating the reproducibility of *B. distachyon* phenotypes, exometabolite profiles, and microbiome assembly within the EcoFAB 2.0 device. The experiment compared the recruitment of the full SynCom vs. one lacking the dominant root colonizer *Paraburkholderia* sp. OAS925 ^11^. To minimize variation required in all laboratories, almost all supplies (e.g., EcoFABs 2.0, seeds, SynCom inoculum, filters) were distributed from the organizing laboratory, and detailed protocols, including annotated videos, were created. Each laboratory measured plant phenotypes and collected samples for 16S rRNA amplicon sequencing and metabolomic analyses by LC-MS/MS. A single laboratory performed all the sequencing and metabolomic analyses to minimize analytical variation. Follow-up *in vitro* assays and comparative genomics were conducted to gain insights into mechanisms leading to *Paraburkholderia* sp. OAS925 dominance. Overall, the study demonstrates consistent plant traits across multiple laboratories and provides publically accessible benchmarking data for other labs to leverage, replicate, and extend this work. In addition, we describe the challenges we encountered in performing this study, thus providing information that can facilitate future microbiome reproducibility studies.

## 2. Results

### 2.1. Standardized protocols achieve EcoFAB 2.0 device sterility

Our main objective was to develop and test methods to reproducibly study plant microbiomes within the sterile EcoFAB 2.0 device (**Fig.1a**). We hypothesized that the inclusion of *Paraburkholderia* sp. OAS925, a dominant *B. distachyon* root colonizer into SynCom ^11^, would reproducibly influence the microbiome assembly, metabolite production, and plant growth across multiple laboratories using the EcoFAB 2.0 device. To test the hypothesis, we deployed the grass *B. distachyon* with a SynCom consisting of 16 or 17 members that was originally developed to span the diversity of bacteria isolated from grass rhizosphere, including representatives from the Actinomycetota, Bacillota, Pseudomonadota, and Bacteroidota phyla (**Fig. 1b**) ^9^. Our study was conducted across 5 laboratories (designated A-E) and consisted of four treatments with 7 biological replicates each (**Fig. 1a**): an axenic (mock-inoculated) sterile plant control, SynCom16-inoculated plants, SynCom17-inoculated plants, and plant-free medium control. Each laboratory followed written protocols and annotated videos, gathered root and unfiltered media samples for 16S rRNA amplicon sequencing, filtered media for metabolomics, measured plant biomass, and performed root scans. At the end of the study, the collected data and samples were sent to the organizing laboratory for sequencing, metabolomics, and data analysis.

**Figure 1:**
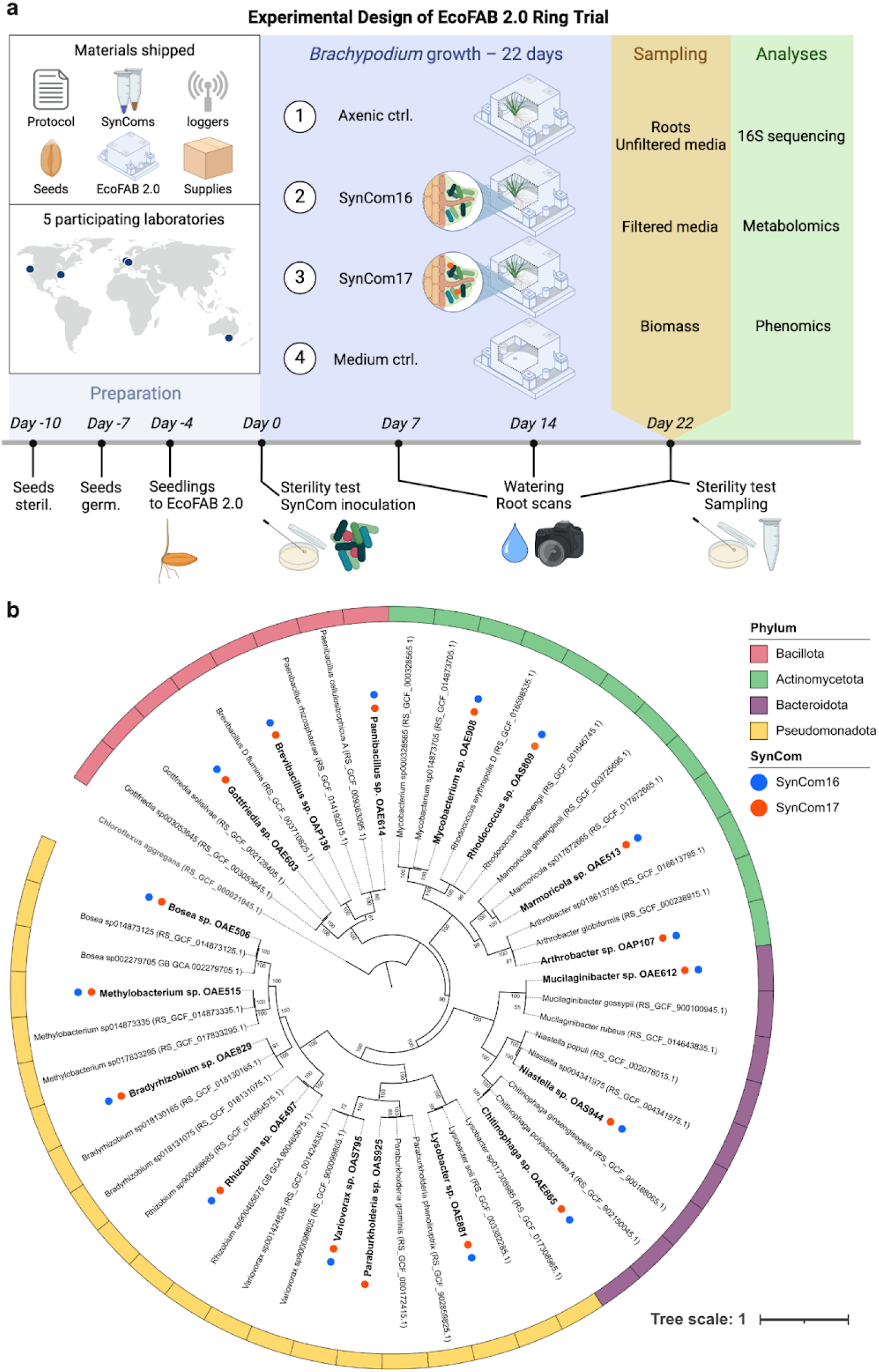
(**a**) Experimental design where five laboratories across three continents conducted the same experiment using shipped materials. These included a detailed protocol (https://dx.doi.org/10.17504/protocols.io.kxygxyydkl8j/v1), SynComs and Mock solution stocks, light and temperature loggers, *Brachypodium distachyon* seeds, EcoFAB 2.0 device parts, and various lab supplies (growth medium, filters, sampling tubes). We inoculated *B. distachyon* plants with either a 16- or 17-member SynCom, with controls being axenic plants and medium-only (Mock-inoculated), *n*=7. We tested sterility and imaged roots at multiple time points. Finally, we quantified plant biomass, analyzed exudate metabolite composition, and measured root and medium microbiomes. (**b**) The phylogenomic tree is based on 120 marker genes, where SynCom members are highlighted in bold, with phylum-level classification shown by colored strips and SynCom membership by circles (SynCom16 in blue, SynCom17 in orange). The 2 closest taxonomic genomes are included, with GenBank accession numbers in parentheses. Nodes with over 50 bootstrap support values from 100 replicates are labeled. *Chloroflexus aggregans* (bold gray) served as an outgroup.

The detailed protocol with embedded annotated videos used by all five laboratories is available via protocols.io (https://dx.doi.org/10.17504/protocols.io.kxygxyydkl8j/v1). The general procedure follows these steps: (i) EcoFAB 2.0 device assembly; (ii) *B. distachyon* seed dehusking, surface sterilization, and stratification at 4 °C for 3 days; (iii) Germination on agar plates for 3 days; (iv) Transfer of seedlings to the EcoFAB 2.0 device for additional 4 days of growth; (v) Sterility test and SynCom inoculation into the EcoFAB 2.0 device; (vi) Water refill and root imaging every seven days; (vii) Sampling and plant harvest at 22 days after inoculation (DAI). Since differences in labware and material can cause experimental variation, the protocol specifies the part numbers used in this study. Organizers provided critical components, including growth chamber dataloggers, in the initial package of non-perishable supplies, while the SynComs and freshly collected seeds were shipped just before the study. Given the time zone differences, it was difficult to synchronize all activities, so each laboratory performed the experiment independently within 1.5 months of each other (**Table S1**). All participants followed data collection templates and image examples.

### 2.2. Protocols resulted in reproducibly sterile conditions

During the study, all participating laboratories tested the sterility of the EcoFABs 2.0 devices at two different time points, imaged plant roots using a flatbed scanner, and quantified root fresh and shoot fresh and dry biomass during harvest. The sterility of uninoculated devices was tested by incubating spent medium on Luria-Bertani (LB) agar plates. Less than 1% (2 out of 210) of all tests showed colony formation (**Fig. 2a)**. Namely, a single colony was observed in one treatment of laboratory D in SynCom17, and multiple colonies for laboratory B in medium-only control (plate had cracked lid).

**Fig. 2:**
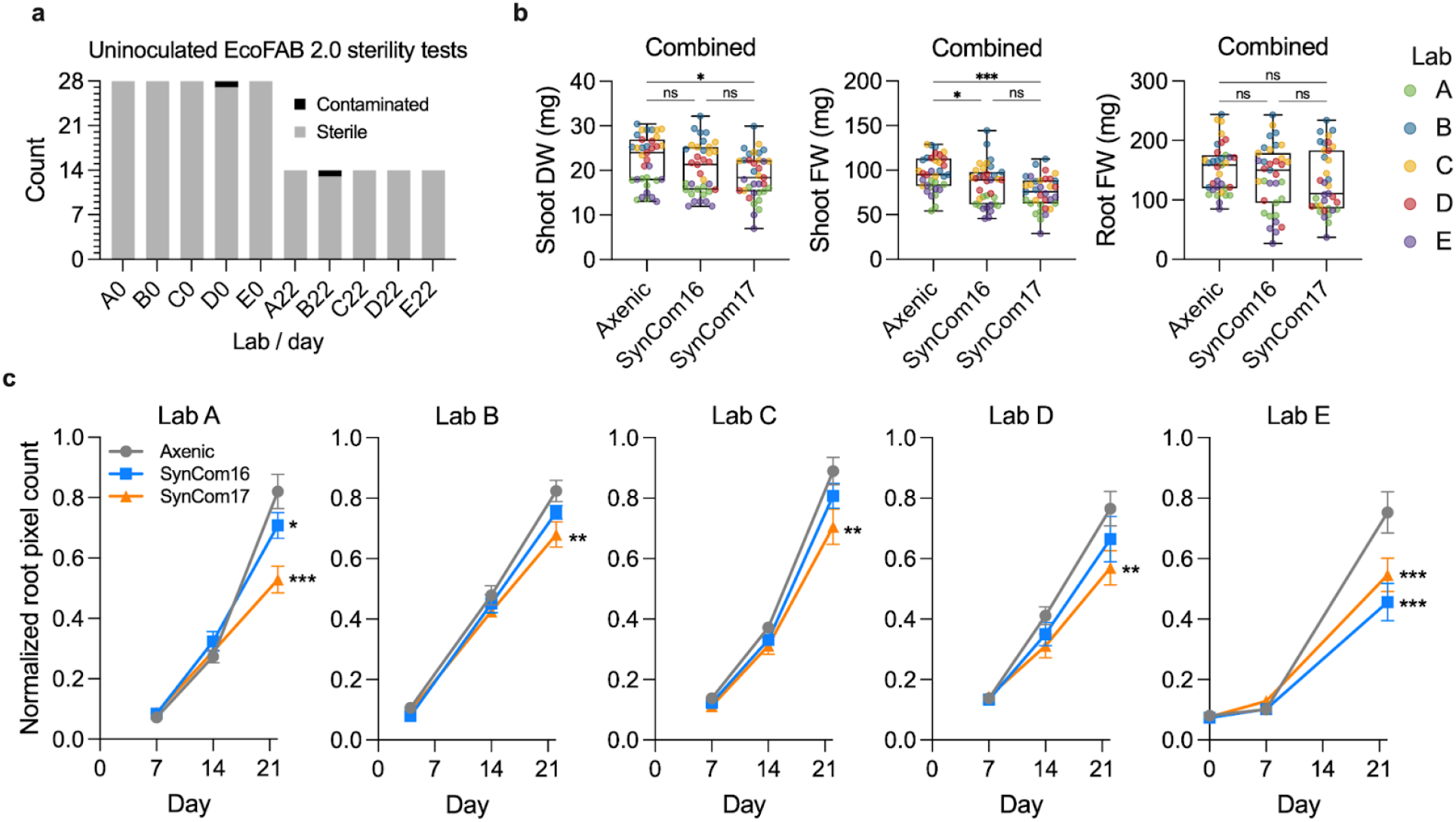
P**l**ant **phenomics and EcoFAB 2.0 device sterility**: (**a**) Sterility of uninoculated EcoFAB 2.0 devices tested across laboratories A-E at day 0 and day 22 after inoculation. The medium from these devices was incubated on LB agar plates for 22 days to observe bacterial colony formation. (**b**) Plant biomass weight combined across all laboratories (Lab A-E in different colors), measured as shoot dry weight, shoot and root fresh weight. One-way ANOVA with Tukey test, *n* = 7, ns *p*>0.5, \**p*<0.05, \*\**p*<0.01, \*\*\**p*<0.001. (**c**) Root system development was analyzed using RhizoNet (Lab B-E) and ImageJ (Lab A). The raw root pixel counts were normalized to the maximum value in each lab. Two-way ANOVA with Dunnet’s test vs. Axenic control, *n* = 7, ns *p*>0.5, \**p*<0.05, \*\**p*<0.01, \*\*\**p*<0.001.

### 2.3. Reproducible plant growth

When plant biomass data were combined across laboratories, we observed a significant decrease in shoot fresh weight and dry weight of plants inoculated with SynCom17 relative to the axenic treatment (**Fig. 2b)**. This said, we did observe some variability between laboratories (**Fig. S1**), which is presumably due to growth chamber differences including light quality (fluorescent vs LED growth lights), light intensity and temperature (**Table S1**). Supporting this, the data loggers revealed variability in measured temperatures (**Fig. S2a)** and photoperiod (**Fig. S2b)**. Image analysis of scanned roots revealed that SynCom17 caused a consistent decrease in root development observed after 14 DAI onwards (**Fig. 2c**).

### 2.4. Reproducible microbiome assembly

SynComs were prepared using optical density (OD_600_) to colony-forming unit (CFU) conversions (**Table S2)** to ensure equal cell numbers (final inoculum 1e5 bacterial cells per plant) and shipped on dry ice to each laboratory as 100x concentrated stocks in 20% glycerol. The cells were resuspended and added to 10-day-old *B. distachyon* seedlings in the EcoFAB 2.0. After 22 days of growth, the roots and media were sampled, shipped back to the organizing laboratory, sequenced, and compared to the original inoculum. For both SynComs (SynCom16 and SynCom17), the community composition at 22 DAI differed from the inoculum (**Fig. 3**). As hypothesized, the root microbiome inoculated with SynCom17 was dominated by *Paraburkholderia* sp. OAS925 across all laboratories (98±0.03% average relative abundance ± SD). In its absence (SynCom16), other isolates showed high relative abundance in the root microbiome with increased variability across laboratories, namely *Rhodococcus* sp. OAS809 (68±33%), *Mycobacterium* sp. OAE908 (14±27%), and *Methylobacterium* sp. OAE515 (15±20%). The most dominant microbial isolates detected in root samples were also typically present in the media samples (**Fig. S3a**). Ordination plots showed clear separations between SynCom16 and SynCom17 microbiomes for both root and media, with generally higher variability between samples for the SynCom16 microbiome (**Fig. S3b)**. There was a minimal contribution of unknown reads in all samples, consistent with the observed sterility of the controls. Furthermore, SynCom17 treatment in laboratory D did not show unknown reads, suggesting that the negative sterility test (**Fig. 2a**) was likely caused by plate contamination.

**Figure 3:**
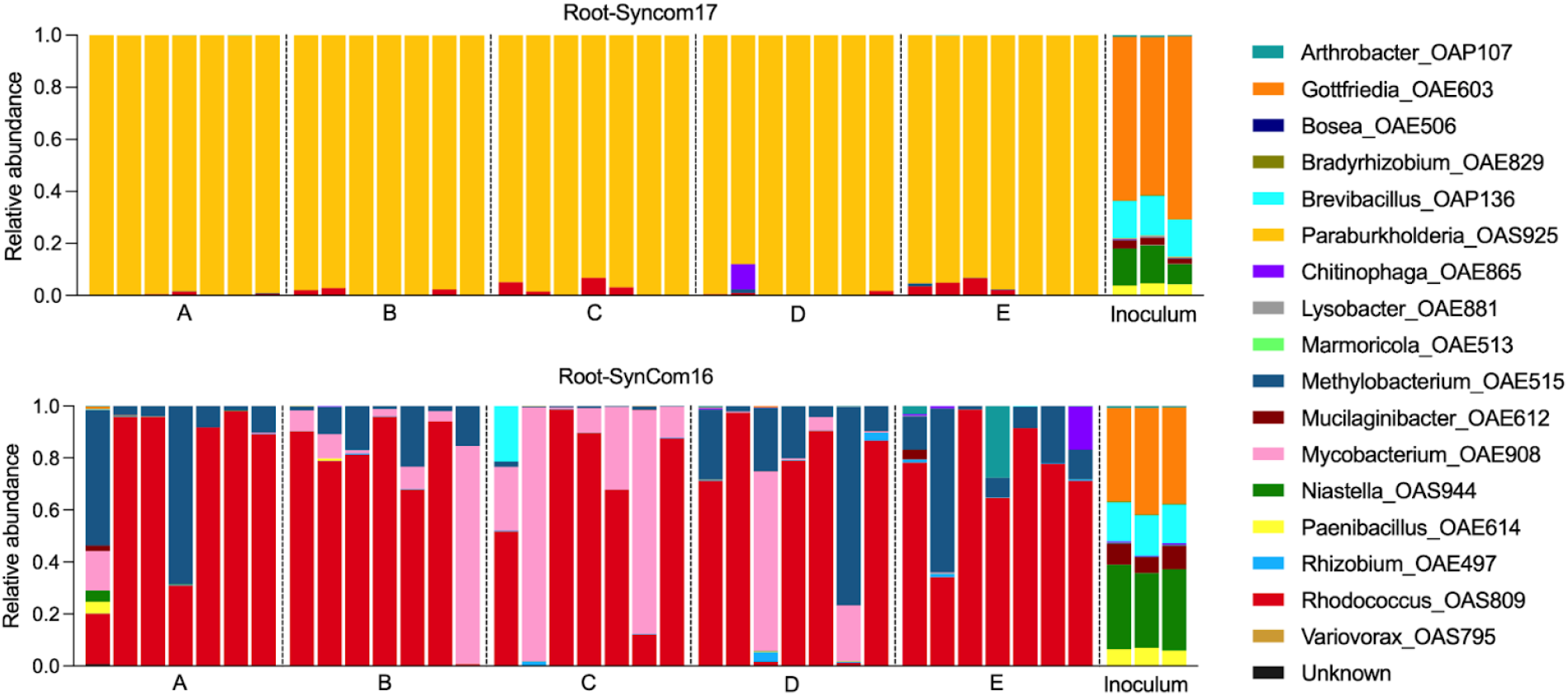
R**o**ot **microbiome**. Microbiome composition of *Brachypodium distachyon* roots and starting inoculum of plants inoculated with SynCom16 or SynCom17 (±*Paraburkholderia* sp. OAS925). Letters indicate different laboratories, with each biological replicate shown (*n*=7). The inoculum shows technical replicates (*n*=3).

### 2.5. Reproducible rhizosphere metabolome

The spent medium from each fabricated ecosystem was filtered and shipped to the organizing laboratory for LC-MS/MS analysis (polar HILIC in positive mode, **Table S3**), followed by targeted and untargeted metabolomics to determine the root exudate composition and metabolite profiles in the presence of different SynComs in the rhizosphere. The targeted analysis identified 60 metabolites spanning diverse metabolite classes (**Table S4**). Hierarchical clustering revealed general clustering by treatment and not laboratory (**Fig. 4**), consistent with the experimental reproducibility observed with plant growth phenotypes and root microbiome composition. Furthermore, the metabolite clustering showed several treatment-dependent metabolite changes. The first large cluster included diverse metabolites increased in the SynCom17 treatment. A second large cluster consisted of metabolites with lower relative concentrations in the SynCom17 treatment, represented mainly by amino acids. A third, much smaller cluster consisting primarily of nucleosides(tides) increased in the SynCom16 or both SynCom treatments. This finding highlights the prominent impact of the community dominated by *Paraburkholderia* sp. OAS925 on modulation of metabolite composition in the rhizosphere. This was further supported by untargeted metabolomics on 833 detected features that showed a clear separation between rhizosphere metabolomes of axenic plants and SynCom17, which was reproducible across all laboratories (**Fig. S4**). These changes may be due to metabolite production or uptake by the microbes or plant roots or the activity of extracellular enzymes ^15–17^.

**Figure 4:**
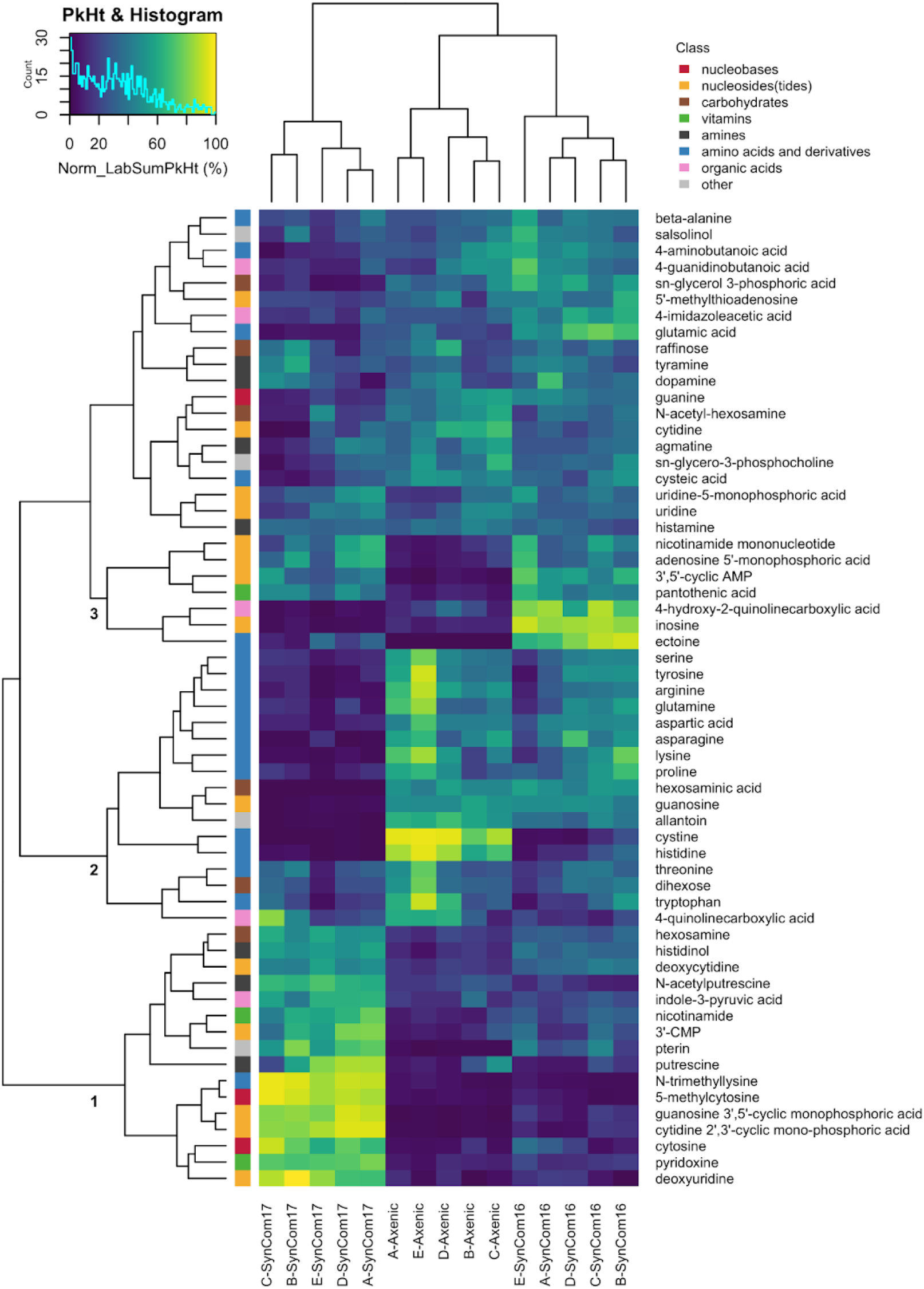
T**a**rgeted **metabolomic analysis of rhizosphere.** We show mean values for each lab/treatment combination (*n*=7), row-normalized to the average sum peak height per lab. Row colors indicate metabolite classes. Cluster 1: Abundant in SynCom17; Cluster 2: Low in SynCom17; Cluster 3: Abundant in SynCom16 or both SynComs.

### 2.6. Colonization by *Paraburkholderia*

Given the reproducible changes in our fabricated ecosystems, including plant growth phenotypes, microbiome structure, and rhizosphere exometabolites in response to *Paraburkholderia* sp. OAS925, we performed additional analyses to gain insights into potential mechanisms explaining its dominance. Comparative genomic analysis shows that the *Paraburkholderia* sp. OAS925 genome (IMG/M Taxon ID: <u>2931840637</u>) uniquely includes acid resistance genes such as glutamate and arginine transporters and decarboxylases (**Fig. S5**) and a gene module coding for a Type 3 Secretion System (T3SS), which was not found in any other member of the SynCom (**Table S5**).

We inoculated *B. distachyon* with a red fluorescent protein (RFP) expressing *Paraburkholderia* sp OAS925 to investigate spatial-temporal root colonization in EcoFAB 2.0. Clear RFP signals were detected at the root tip and in the maturation zone at 1 DAI, with increased biofilm formation observed at 3 DAI (**Fig. S6**). We noted both sessile colonies on the rhizoplane (**Video S1**) and active swimming surrounding root cells (**Video S2**).

The biofilm formation and motility observed during microscopy motivated the follow-up *in vitro* assays to further assess these characteristics across isolates. *Paraburkholderia* sp. OAS925 exhibited the sixth-highest biofilm formation but the highest growth on a liquid Northen Lab Defined Medium (NLDM) (**Fig. S7**), highlighting its potential to outgrow competitors when cultured with common soil metabolites in the NLDM ^18^.

Swimming motility assays on soft agar revealed that *Paraburkholderia* sp. OAS925 had the highest motility within the first 24 hours (**Fig. S8a)**. Additionally, compared to other isolates with similar motility phenotypes (**Fig. S8b**), it maintained fast swimming in acidic conditions (**Fig. S8c**) in the range of the hydroponic medium (pH 5.5-6.0 at the start of the experiment). KEGG mapping (<u>map02040</u>) showed the presence of flagellar assembly genes, suggesting that the observed motility is due to flagella.

## 3. Discussion

There is an urgent need to move towards replicable experimental systems to address common difficulties in reproducing microbiome experiments ^2,19^. Here, we report what, to our knowledge, is the first multi-laboratory microbiome reproducibility study. We constructed fabricated ecosystems using two SynComs, the model plant *B. distachyon* Bd21-3, and the sterile EcoFAB 2.0 devices. These, in combination with written protocols and annotated videos, resulted in reproducible plant growth phenotypes, host microbiomes, and exometabolomes across five laboratories spanning three continents. Specifically, SynCom17, which contained the dominating bacteria *Paraburkholderia* sp. OAS925 reduces root growth rate and fresh/dry shoot biomass of *B. distachyon*. This finding is consistent with a previous study showing *Paraburkholderia* sp. OAS925 dominance of the *B. distachyon* root and rhizosphere microbiota and decreased fresh root biomass (21 DAI for plants inoculated 8 days after germination) ^11^. Furthermore, similar results were observed for another grass, *Avena barbata*, grown in its native soil, where members of the order *Burkholderiales* were the most active bacteria in the rhizosphere based on carbohydrate depolymerization ^20^.

Soil pH, organic carbon availability, oxygen levels and redox status are key factors influencing microbial community composition ^21^. Our study suggests that *Paraburkholderia*’s dominance in the rhizosphere is due to its genetic and functional capabilities interacting with those factors. The high motility of *Paraburkholderia* sp. OAS925 in acidic environments (**Fig. S8c**), such as the rhizosphere, might facilitate quick colonization of ecological niches and affect community assembly ^22–24^, while its ability to utilize amino acids like arginine, glutamine, and glutamate (**Fig. 4**) provides a possible mechanism for cytoplasm de-acidification to maintain motility-enabling transmembrane proton gradient ^25,26^. These results align with a previous study showing that *Pseudomonas simiae* genes involved in motility, carbohydrate metabolism, cell wall biosynthesis, and amino acid transport aid in *Arabidopsis* root colonization ^27^. Interestingly, in SynCom16, *Rhodococcus* often dominates on roots (**Fig. 3**) and shares fast growth (**Fig. S7**) and high motility (**Fig. 8a**) with *Paraburkholderia*.

The observed invasive colonization by *Paraburkholderia* (**Fig. S6, Video S1 and Video S2**) might disrupt plant nutrient homeostasis, as the root microbiome plays a crucial role in forming root diffusion barriers and maintaining plant mineral nutrient balance ^5^, which could explain the observed decrease in root biomass (**Fig. 2a**). Furthermore, the observation of a T3SS (**Table S5**) is consistent with previous findings in *Paraburkholderia* genomes and has been shown to play a role in root colonization and virulence. Future studies should investigate the role of T3SS in the dominance of *Paraburkholderia* sp. OAS925 in SynCom17 treatments and the associated plant biomass decrease. Additionally, future testing if *Paraburkholderia* sp. OAS925 causes detrimental effects on plant growth in mono-association could indicate whether it is an opportunistic root pathogen whose activity is either insufficiently suppressed by other SynCom members or requires specific strains in the natural root microbiota for suppression.

By organizing this ring trial, we learned valuable lessons that can be useful for future studies (**Fig. 5**). First, it is important to perform pilot studies to optimize methods before initiating any multi-laboratory study. Long-distance sample and inoculum shipping posed challenges, especially given unpredictable long delays in customs and potential thawing due to dry ice sublimation ^28^. Microorganism shipments require engagement with shippers and familiarity with country-specific import/export legal regulations. We also observed variability in plant biomass (**Fig. 2b**), which could be attributed to differences, especially in light and temperature (**Fig. S2**) between the growth chambers used in each laboratory (**Table S1**). Ideally, the same equipment would be used, with a real-time readout of environmental conditions, although this would significantly increase the cost of the study.

**Figure 5:**
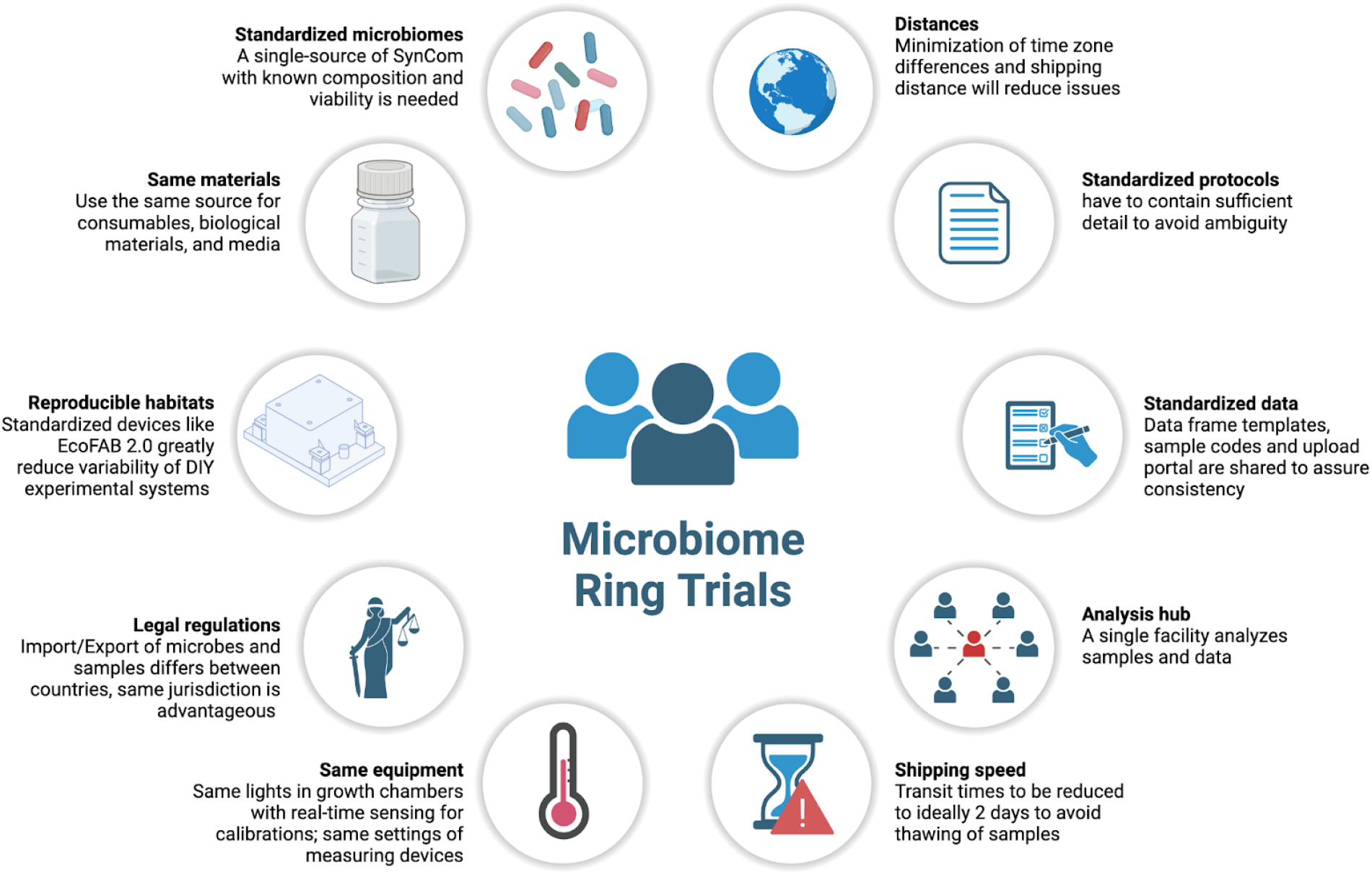
C**o**nsiderations **and Challenges for Reproducible Microbiome Studies**. This figure summarizes key factors and lessons for standardizing microbiome experiments. It highlights the essential elements for organizing reproducible studies and offers insights into the organization of future microbiome multi-lab ring trials.

Despite our detailed protocols and annotated videos, several challenges remain to replicate microbiome studies, underscoring the importance of using the data from this study to benchmark future studies. We recommend using the comment section on protocols.io for ongoing refinement and clarification, allowing the procedure to evolve as a living document ^29^. To provide FAIR (Findable, Accessible, Interoperable, and Reusable) data access and enable others to use these data for benchmarking, integration, and extension, all data from this study are available via the National Microbiome Data Collaborative (NMDC) project page (https://data.microbiomedata.org/details/study/nmdc:sty-11-ev70y104)^30^. The EcoFAB 2.0 devices, *B. distachyon* Bd21-3 plant line, and metabolomics methods used in this study are currently freely available via Joint Genome Institute (JGI) User Programs, while the 16S rRNA sequencing is readily available via commercial and academic sequencing centers. Although the relative abundance of organisms should ideally not correlate with the sequencing facility, sample handling, DNA extraction, and bioinformatics can significantly impact results, underscoring the need to consider protocols when making comparisons ^31^. Another challenge we see is that the strains of our SynCom are currently available as individual strains, so batch variation would be reduced if culture collections or private companies provided ready-to-use SynCom mixtures.

This study demonstrates that multiple geographically dispersed laboratories can reproduce SynCom-driven changes in plant phenotypes, community assembly, and exometabolite profiles. This was a challenging yet essential step in the vision outlined by Zengler et al. ^1^ to verify the reproducibility of experimental systems and protocols, which enable scientists to replicate and build on each others’ work. We see several ways these methods can help advance the field: first, scientists can replicate the study and compare their results against those reported here before extending the findings with additional modifications (e.g., adding phages, fungi, engineered strains, different hosts, new devices, etc.). Second, scientists can generate experimental data through replication and benchmarking, enabling integrative computational analyses that control laboratory-specific effects. Providing FAIR data and accompanying metadata and protocols, as done here, will be an essential step in achieving this vision. Such efforts would greatly enhance the application of machine learning to make generalizable discoveries drawn from multiple studies, ultimately leading to understanding microbial processes in complex natural environments.

## 4. Material and Methods

### 4.1. Preparation of synthetic bacterial communities and their distribution

The bacterial isolates were obtained initially from the rhizosphere of a single switchgrass plant and are available from DSMZ ^9^. These isolates were kept as glycerol stocks. Before initiating work, the 16S gene (27F - 1492R) for each of the 17 isolates was sequenced to verify the identity and confirm purity. Then, each isolate was streaked on an agar plate with an isolate-specific medium (**Table S2**), and a single colony was inoculated into liquid culture. After 2–8 days, depending on the growth of the strain, the cultures were pelleted by centrifugation at 5000 g and then washed with ½ strength Murashige and Skoog (MS) basal salts medium.

The washed cultures were then used to create SynCom stock solutions. First, OD_600_ was measured, and then isolates were combined using the CFU to OD conversion table (**Table S2**) to equal CFU (1:1) of each strain at 2e7 cells/ml into a solution of 20% glycerol, which was shown to be efficient for community cryopreservation ^9^. We also prepared a mock solution of 20% glycerol in ½ MS basal salts. Then, 100 µl of each solution was aliquoted into 1.5 ml Eppendorf tubes and stored at -80°C until shipped. The participants diluted the SynCom and mock stock solutions 100-fold before use. We assumed a general 50% cell survival rate during freezing-thawing. Therefore, the final theoretical CFU of each strain was 1e5/plant. The CFU to OD conversion factors (**Table S2**) were established by plating sequential dilutions of washed cultures with known OD_600_ followed by colony enumeration and hemocytometer for the *Gottfriedia* sp. OAE603.

The inoculums were distributed among five participants: Lawrence Berkeley National Laboratory, USA (organizer); the University of Melbourne, Australia; the University of North Carolina at Chapel Hill, USA; Forschungszentrum Jülich, Germany; and the Max Planck Institute for Plant Breeding Research, Germany. The shipping was optimized for speed by testing different vendors by shipping dummy tubes before proceeding with the actual shipment of real SynCom samples. All paperwork for the sample shipment was obtained ahead of time, and a content declaration form with a cover letter from the receiving laboratory was included. We encountered diverse and sometimes ambiguous regulations for shipping living microbiomes across international borders. Most countries prohibit the import of listed pathogens. Therefore, it is advantageous to identify the closest known phylogenetic species for each bacterial isolate and list them on the shipment declaration to avoid delays at customs. Shipment of SynComs to Germany was not explicitly regulated and was labeled BSL1. For both shipments to Germany, we used FedEx International Priority with 8.2 kg of dry ice and shipped the packages as a delivered-at-place (DAP) with a proforma invoice from the consignor containing freight charges and the value of the goods. The SynCom shipment to Australia required a permit to import conditionally non-prohibited goods under the Biosecurity Act 2015 Section 179 (1). For the shipment, we used Aeronet Worldwide with 22.68 kg of dry ice, and the company guaranteed a refill of dry ice during shipment, which took 9 days to deliver. The SynCom shipment to North Carolina was not regulated, and we used FedEx Standard Overnight with 9 kg of dry ice (1 day in transit). All deliveries arrived frozen.

### 4.2. Experimental setup and plant growth conditions

Participating laboratories assembled EcoFAB 2.0 devices and followed experimental procedures described in the protocol (https://dx.doi.org/10.17504/protocols.io.kxygxyydkl8j/v1). In summary, seeds of *B. distachyon* Bd21-3 ^32^ were surface-sterilized by washing in 70% ethanol for 30 s, followed by 5 min wash in 6% sodium hypochlorite solution. Afterward, the seeds were washed 5 times with sterile milli-Q water. Seeds were then plated on plates with ½ MS basal salts and 1.5 % (w/v) phytoagar. After stratification for 3 days in the dark at 4°C, plates with seeds were moved to the growth chamber with a 14 h photoperiod at 26°C and 10 h dark at 20°C. The photosynthetic photon flux density (PPFD) was 110-140 µmol/m^2^/s across laboratories. If tunable, humidity was set to 70%.

Each lab placed two data loggers, HOBO MX2202 (Bluetooth-readout) and HOBO UA-002-64 (coupler-readout), to track illuminance (lux) and temperature. Due to variability in logger placement and resulting lux readings, illuminance was solely used to confirm night period duration, not as a proxy for photosynthetically active radiation ^33^. After 3 days, germinated seeds were aseptically transferred into autoclave-sterilized EcoFAB 2.0 filled with 9 mL of ½ MS basal salts medium adjusted to pH 5.5-6.0 with KOH. The medium was filter-sterilized using 0.2 µm PES membrane 1L filtration units. The EcoFABs 2.0 devices were then placed back in the chamber following the germination settings. After 4 days in EcoFAB 2.0, plants were inoculated. Briefly, 100 µl of stocks of Mock, SynCom16, and SynCom17 solutions with 20% glycerol shipped to the laboratories were resuspended to 10 ml in sterile ½ MS basal salts solution. Then, 1 ml of treatment solution was added to 9 ml of medium within the EcoFAB 2.0 devices. The treatment groups included: i) Mock-inoculated axenic plant control, ii) SynCom16-inoculated plants, iii) SynCom17-inoculated plants, and iv) Mock-inoculated technical control (plant-free with medium only). Therefore, all EcoFAB 2.0 devices had a final glycerol concentration of 0.02%. Sterility of the uninoculated EcoFABs 2.0 devices was tested at day 0 just before inoculation for all devices and at day 22 after inoculation for plant axenic and technical controls by plating 50 µl of the spent medium on LB agar plates and incubating for 22 days at room temperature to test for formation of colonies due to contamination. At 22 DAI, plants were harvested and media sampled.

## 5. Plant phenotyping

To automate analysis of root development utilizing root scans, we deployed RhizoNet, a deep learning computational workflow designed to precisely segment plant roots that is particularly well suited for the tangled roots imaged through the bottom of the EcoFAB 2.0 devices^34^. We encountered challenges during the analysis of scans from Lab A using RhizoNet due to condensation and reflections during scanning that resulted in low contrast. Therefore, the roots for Lab A were analyzed with ImageJ V1.54 ^35^ coupled with the SmartRoot plug-in V4.21 ^36^, followed by calculating the total root length for each scan by summing the length of the first-order roots. The raw root measurements from both methods were normalized to the maximum value across all time points for each lab. The root and shoot were separated, and then the fresh weight biomass of roots and shoots was measured. The roots were then frozen for subsequent microbiome analyses. The shoots were frozen, then lyophilized, and dry weight was recorded.

### 5.1. Sample collection and shipment

Sample collection and initial processing were performed independently in each lab. To determine microbiome composition in the growth medium, 50 µl of the unfiltered growth medium was collected into 1.5 mL Eppendorf tubes for 16S rRNA amplicon sequencing. Then, to determine root exudate composition, all remaining spent medium was filtered via a 0.2 µm PES syringe filter for metabolomics. Exudates for metabolomics were collected in 15 mL polypropylene conical tubes, all from the same product line but with either HDPE plug caps for laboratories B-E (VWR 93000-026) or with HDPE flat caps containing thermoplastic elastomeric sealing ring for lab A (VWR 21008-103), due to product availability. Exudates were collected by each lab and stored at -80 °C before shipment. The EcoFAB 2.0 device was then opened, and the root and shoot separated. The fresh root was placed in a pre-weighed 2 ml Eppendorf tube, the fresh weight of the roots was determined, and the roots were frozen for 16S rRNA amplicon sequencing to determine root microbiome composition. Plant shoots were frozen and then lyophilized to complete dryness, and then dry shoot biomass was determined.

The filtered medium (for LC-MS/MS), unfiltered medium, and frozen roots (for amplicon sequencing) were shipped to Lawrence Berkeley National Laboratory, California, USA. The sample shipments are required to be timely and temperature-sensitive. All samples were shipped on dry ice with additional gel packs. The pendant data loggers were shipped separately at room temperature to download data. To import intact frozen *B. distachyon* roots to the USA, we have obtained a Controlled Import Permit to Import Restricted or Not Authorized Plant Material Regulated by 7 CFR 319.6 (PPQ Form 588) for each country of origin. Each sample import shipment was accompanied by supplier declaration and Toxic Substance Control Act (TSCA) Certification. We imported samples from Germany by DHL Medical Express (2 days in transit) on 10 kg of dry ice. For shipment from Australia, we used Cryopdp (5 days in transit) on 24 kg dry ice, and we requested that the vendor refill the dry ice during shipment. The shipping between North Carolina (Orange County) and California did not require specific permits as directed by the California Department of Food and Agriculture. For the shipment between North Carolina and California, we utilized FedEx Priority Overnight on 9kg dry ice (1 day in transit). All samples arrived frozen.

### 5.2. Root exudate metabolomics and data analysis

At Lawrence Berkeley National Laboratory, samples were removed from the freezer and dried by lyophilization (Labconco, Kansas City, MO). Tube caps from lab A with a gasket (VWR 21008-103) were replaced with caps used by all other laboratories (VWR 93000-026) to maintain consistency and avoid material contamination differences during extraction. Empty tubes (VWR 21008-103 with caps from VWR 93000-026 and tubes and caps from 93000-026) were used as extraction controls. All samples were kept on dry ice during the following extraction process. Solvents were chilled at -20°C before extracting. The dried material was suspended in 1mL methanol (MX0486 Omnisolv LCMS grade, Sigma), vortexed 2 x 10 s, transferred to a 2mL microcentrifuge polypropylene snap cap tube (022431048, Eppendorf); the 15mL tube was washed with an additional 0.5mL methanol by pipetting up and down to collect residual dried material and combined in the 2mL tube. Samples were then bath sonicated (97043-944, VWR) in ice water for 15 minutes and then centrifuged at 10,000g for 5 min at 10°C to pellet insoluble material. Supernatants were transferred to a second set of 2mL tubes and then dried by vacuum concentration overnight (SpeedVac system with RVT5105-115 concentrator and SPD130DLX centrifuge, Thermo). The following morning, samples were removed from the SpeedVac centrifuge and resuspended in 150 µL methanol containing internal standard mix (**Table S3**). Samples were vortexed 2 x 10 s and centrifuged at 10,000g for 5 min at 10°C. Then, to filter the samples, the supernatant was transferred to centrifugal polypropylene tubes with 0.22µm hydrophilic PVDF filters (UFC30GV Ultrafree-MC Centrifugal Filter, Millipore) and centrifuged at the same settings as above. Filtrates were then collected in amber glass vials with 300uL inserts (5188-6592, Agilent) and immediately capped with polypropylene screw caps with PTFE/silicone septa (5185-5820, Agilent).

Samples were analyzed using LC-MS/MS. Briefly, polar metabolites were separated using hydrophilic liquid interaction chromatography (InfinityLab Poroshell 120 HILIC-Z, 2.1 x 150 mm, 2.7 µm column, 683775-924, Agilent) on an Agilent 1290 HPLC system followed by detection on a Thermo Orbitrap Exploris 120 Mass Spectrometer equipped with an H-ESI source probe using data-dependent MS2 acquisition to select the top two most intense ions not fragmented in the previous 7 s. Samples were injected in positive mode, with methanol blanks injected between each sample; internal and external controls were used for quality control. LC-MS/MS parameters are described in **Table S3**. Thermo raw files were converted to mzML format using ThermoRawFileParser ^37^.

For untargeted metabolomics, the mzML files were processed via MZMine 3.0 ^38^ to create lists of features and MGF MS2 container files using a custom batch process (**File S1**). The features with MS2 spectrum were then annotated in GNPS2 (Global Natural Products Social Molecular Networking) using spectral metabolite libraries ^39^. The total number of features was then filtered to include features with MS2, RT > 0.6 min, and maximum exudate sample peak height > 10x of extraction and technical control samples (for fold-change calculations, +1 was added to the numerator and denominator values). This resulted in a total of 833 features across all samples and laboratories.

For targeted metabolomics, metabolites were identified (level 1) (**Table S4**) by analyzing the data with an in-house library of m/z, RT, and MS2 fragmentation information from authentic reference standards using Metabolite Atlas (https://github.com/biorack/metatlas) ^40,41^. Only metabolites with a maximum exudate sample peak height > 3x of extraction and technical controls were included. The identified 70 metabolites were manually classified using the PubChem Classification Browser ^42^.

### 5.3. Microbiome composition analysis by 16S rRNA amplicon sequencing

DNA was extracted from the ground roots and media samples using the DNeasy PowerSoil Pro Kit (Qiagen), following the manufacturer’s instructions with minor modifications. Ground roots were suspended in the resuspension buffer, transferred to bead-beating tubes, and frozen at -80°C before extraction. All samples were then thawed at 60°C, and bead beating was conducted using the FastPrep-24 sample preparation system for 30 s at setting 5.0 (MP Biomedicals). The elution buffer was heated to 60°C before use. Samples were extracted in batches, and a water-only sample was included for each batch as a negative control. For time zero data, 3 glycerol stocks for each 16 and 17-member communities were performed along with a glycerol-only negative control.

PCR amplification of V4 amplicons was performed in two steps. Library amplicons were generated using the Illumina i7 and i5 index/adapter sequences with V4 priming sequences 515F (GTGYCAGCMGCCGCGGTAA) ^43^ and 806R (GGACTACNVGGGTWTCTAAT) ^44^ using a two-step process. First, amplification was performed on all samples, including negative controls, using pooled primers on a Bio-RAD CFX 384 Real-Time PCR detection system using the QuantiNova SYBR Green PCR kit (Qiagen) in 10 µl reactions with primers supplied at 4 µM and mitochondrial and chloroplast PNA blockers (PNA Bio) supplied at 1.25 µM. Amplification was initiated at 95°C for 3 min, followed by the following cycle: 95°C for 8s, 78°C for 10s, 54°C for 5s, and 60°C for 30s, followed by fluorescence measurement. Root and media samples were evaluated separately, along with negative and positive controls, to identify the number of cycles where most samples reached the late-exponential phase. Two libraries were prepared, one with media and one with root samples. Root samples were amplified for 22 cycles and media samples for 30 cycles with at least 1 replicate. Samples that went well into the plateau phase were diluted, and rerun, and more replicates were performed on low-amplifying samples. For time-zero glycerol stock samples, samples were prepared for each library preparation and were diluted accordingly based on the number of amplification cycles.

Libraries were purified at least twice before sequencing to remove excess primers. First, individual reactions were purified using the Mag-Bind TotalPure NGS beads (0.8X) following the manufacturer’s instructions, and targets were quantified using the QuantiFluor dsDNA System (Promega). Libraries were then pooled (i.e., root and media samples were pooled separately) to get equal concentrations of each target, and the pooled mixture was purified at least once more using the Mag-Bind kit.

MiSeq reads were processed using Usearch (v11.0.667) ^45^. Initial read preparation was performed using the ‘fastq_mergepairs, fastx_truncate, and fastq_filter’ commands to merge, trim, and remove short sequences. Then, an initial OTU table was generated with the ‘fastx_uniques, cluster_otus, and otutab’ functions. To assign these OTUs to the SynCom members, the ‘annot’ function was used with V4 reference sequences for the SynCom bacteria.

### 5.4. Biofilm formation assays for bacterial isolates

The crystal violet assay for biofilm formation was modified from the previously published method ^46^. In summary, isolates were grown in R2A, washed, and resuspended in a 30 mM phosphate buffer. They were inoculated into the screening plates (90 µL of NLDM medium^18^ ) at a 1:10 (v/v) ratio to achieve a final volume of 100 µL (initial OD_600_ of 0.02) and incubated statically at 30°C incubator for 3 days (*n*=4–5). Post incubation, the supernatant was discarded, and each well was washed thrice with MilliQ water and air-dried. 125 µL of a 0.1% crystal violet solution (0.1% v/v crystal violet, 1% v/v methanol, and 1% v/v isopropanol in MilliQ water) was added to each well, followed by a 30-minute room temperature incubation. After discarding the staining solution, wells were rinsed thrice with MilliQ water. The biofilms were then destained with 125 µL of a 30% acetic acid solution and incubated at room temperature for 30–60 minutes. OD_550_ of the destaining solution was measured for quantification.

### 5.5. Genomic and phylogenetic analysis of bacterial isolates

All strains were grown in R2A except *Bradyrhizobium* sp. OAE829, which was grown in 1/10 strength R2A. After collecting pellets, we extracted high molecular weight genomic DNA with the Monarch HMW DNA Extraction Kit (New England Biolabs) or the MasterPure Complete DNA Purification Kit (Lucigen). The genomic DNA was submitted to the Joint Genome Institute for long-read sequencing.

The whole genome sequencing for 16 isolates was done using PacBio Sequel II, Award DOI <u>10.46936/10.25585/60001370</u>, while *Mycobacterium* sp. OAE908 was sequenced with Illumina, Award DOI <u>10.46936/10.25585/60001258</u>. Genomes were stored in the Genomes OnLine Database (GOLD) ^47^, followed by submission to the Integrated Microbial Genomes and Microbiomes (IMG/M) (https://img.jgi.doe.gov/) for annotation^48^. The annotated genomes can be accessed via IMG/M under the listed Taxon ID or GOLD Project ID (**Table S2**): [*Arthrobacter* sp. OAP107](https://img.jgi.doe.gov/cgi-bin/m/main.cgi?section=TaxonDetail&page=taxonDetail&taxon_oid=2931867202) (2931867202, Gp0588953), [*Gottfriedia* sp. OAE603](https://img.jgi.doe.gov/cgi-bin/m/main.cgi?section=TaxonDetail&page=taxonDetail&taxon_oid=2931797537) (2931797537, Gp0588949, formerly known as *Bacillus* sp. OAE603 ^11^), [*Bosea* sp. OAE506](https://img.jgi.doe.gov/cgi-bin/m/main.cgi?section=TaxonDetail&page=taxonDetail&taxon_oid=2931782253) (2931782253, Gp0589672), *<u>Bradyrhizobium</u>* <u>sp. OAE829</u> (2931808876, Gp0589676), *<u>Brevibacillus</u>* <u>sp. OAP136</u> (2931855177, Gp0588951), *<u>Paraburkholderia</u>* <u>sp. OAS925</u> (2931840637, Gp0589681, formerly known as *Burkholderia* sp. OAS925 ^9^), *<u>Chitinophaga</u>* <u>sp. OAE865</u> (2931817136, Gp0589677), *<u>Lysobacter</u>* <u>sp. OAE881</u> (2931823763, Gp0589678), *<u>Marmoricola</u>* <u>sp. OAE513</u> (2931787146, Gp0589673), *<u>Methylobacterium</u>* <u>sp. OAE515</u> (2931791092, Gp0589674), *<u>Mucilaginibacter</u>* <u>sp. OAE612</u> (2931861231, Gp0588952), *<u>Mycobacterium</u>* <u>sp. OAE908</u> (2852593896, Gp0440934), *<u>Niastella</u>* <u>sp. OAS944</u> (2931847253, Gp0589682, listed as *Chitinophagaceae* sp. OAS944), *<u>Paenibacillus</u>* <u>sp. OAE614</u> (2931801854, Gp0589675), *<u>Rhizobium</u>* <u>sp. OAE497</u> (2931775946, Gp0589671), *<u>Rhodococcus</u>* <u>sp. OAS809</u> (2931833612, Gp0589680), *<u>Variovorax</u>* <u>sp. OAS795</u> (2931827682, Gp0588950).

Next, we conducted comparative genomics. The genome statistics and the abundance of protein-coding genes connected to KEGG pathways for individual isolates were obtained from IMG/M. Based on the genes involved in acid resistance of *E. coli* that include Glutamine/Glutamate and Arginine membrane transporters (GadC and AdiC), glutaminase (YbaS), and decarboxylases (GadA, GadB, and AdiA) ^25^, we searched for genes annotated with similar functions (GltIJKL, HisPMQ-ArgT, GlnHPQ, glsA, GAD, AdiA) in the isolate genomes in IMG/M. For an overall comparison between genomes, we used the “Statistical Analysis” tool from IMG ^49^ to compare the coverage of KEGG modules, i.e., the number of genes of each module identified per genome, between *Paraburkholderia* sp. OAS925 on one side and the 16 other genomes on the other side. KEGG modules coverage were compared between the two groups using Fisher’s exact test, and modules with a corrected *p*-value < 10^-05^ were manually inspected for a potential link to plant root colonization.

A phylogenomic tree depicting 17 SynCom members was constructed by employing the GTDB-tk workflow ^50^, which incorporates 120 marker proteins to obtain the multiple sequence alignment, which was subsequently used to generate the tree using Fasttree ^51^. Two genomes that were taxonomically closest to the 17 members in the resulting tree were selected to make this tree, which was visualized with Interactive Tree Of Life (iTOL) ^52^. Genbank accession numbers are provided in parentheses, with the 17 SynCom members highlighted in bold within the tree. The NCBI phylum-level taxonomic classification is indicated for each member ^53^. Bootstrap values, derived from 100 replicates, are displayed for nodes with over 50 bootstrap support values. *Chloroflexus aggregans* (gray and bolded) was used as an outgroup to root the tree.

### 5.6. Fluorescent microscopy

The pGinger plasmid 23100 containing the RFP gene under the kanamycin (Kan) resistance marker was introduced into *Paraburkholderia* sp. OAS925, as described previously ^54^. Briefly, 1mL of *Paraburkholderia* sp. OAS925 grown overnight at 30°C in R2A medium was mixed with 1mL of *E. coli* S17 dapE- harboring the pGinger plasmid grown overnight at 37° C on LB medium with Kan at 50 µg/mL and diaminopimelic acid (DAP) at 300 µM. The mixture was pelleted for 1 min at 10000 g and then resuspended in 100 µL water with 300 µM DAP. This mixture was then placed onto an R2A agar plate and incubated overnight at 30° C. The bacterial mix was then scraped, resuspended in water, and plated on R2A with Kan 50 µg/mL. Transconjugants were verified via fluorescent microscopy and colony PCR.

The *Paraburkholderia* sp. OAS925, expressing red fluorescent protein (RFP), was cultured in liquid 1xR2A broth supplemented with 20 mg/L of kanamycin to maintain selective pressure. The culture was grown in a 7 ml volume within a culture tube, shaken at 200 rpm at 27°C in the dark at a 40° angle for aeration. After 24 hours, the culture reached an optical density (OD) of 0.743; the cells were then harvested by centrifugation at 4000 g for 10 minutes. The supernatant was discarded, and the resulting pellet was resuspended in ½ MS basal salts. The washed culture was used to inoculate 3-day-old plants in EcoFABs 2.0 at a starting OD of 0.01. The root systems of the plants in the EcoFAB 2.0 devices were scanned using a flatbed scanner to capture root architecture and to indicate locations for microscopy at 1 and 3 days after inoculation (DAI). The EVOS M5000 imaging system (Thermo Fisher) was used for inverted microscopy by directly placing the EcoFAB 2.0 device into the microscopy platform. In most cases, microscopy images were created by merging txRED and bright field microscopy. Uninoculated plants served as controls for autofluorescence (GFP and TxRed).

### 5.7. Motility assays

Precultures were grown in culture tubes with 8 ml of compatible liquid medium shaken at 200 rpm, at 27°C in the dark, for 5–8 days. The swimming motility was tested by observing colony spreading on plates with nutrient-rich R2A soft agar at 0.3% (w/v) ^55^. First, we tested motility for all 17 isolates at pH 7.2, followed by motility testing for *Paraburkholderia*, *Gottfriedia*, or *Brevibacillus* at pH 4, 5, 6, 7, 8, or 9, adjusted with 1M HCl or 0.5M NaOH. Each plate containing 30 ml of the solidified medium on Petri dishes (Ø=10cm) was inoculated with 5µl of well-grown culture at the center (*n*=3). Plates were incubated in the dark at 27°C, and the motility ring diameter was measured after 24 and 45 hours.

### 5.8. Data sharing, statistical analyses, software, and data visualization

The participating laboratories uploaded plant biomass data, root scans, and photos into a shared Google folder with pre-defined structured directories and Excel spreadsheets. The heat maps were generated with RStudio version 4.0.5 using heatmap.2 in the ggplots package ^56^. GraphPad Prism 10 version 10.2.3 generated all other plots and statistical analyses. Biorender.com was used to create graphical overviews. Microsoft Excel version 16.78.3 was used to store and manipulate data frames.

## 6. Author contributions

T.R.N., J.P.V., and V.N. conceptualized and designed the study. V.N. finalized methods, managed the study, conducted experiments with help from Y.D., analyzed data, created figures, and drafted the manuscript with input from T.R.N. P.F.A. assembled SynComs and conducted microbiome analyses. J.C., C.F., J.M.K., and K.W. conducted experiments in participating laboratories under the supervision of B.A., J.L.D., P.S.L., and M.W. E.K. conducted ImageJ analysis and drafted methods with input from B.L. S.K. conducted LC-MS extractions and analysis and validated the results. C.D. and M.Z. conducted 16S rRNA amplicon sequencing. S.R., T.K.O., and A.M.D. sequenced and analyzed bacterial genomes. Z.S. and D.U. analyzed root phenotypes with RhizoNet. C.A.A. provided a bacterium for fluorescent microscopy. A.Y. conducted phylogenetic analysis, and M.C. conducted biofilm assays under the supervision of R.C. K.Z. provided funding and reviewed the manuscript. All authors provided comments and approved the final version.

## Acknowledgments

We would like to thank Julio Corral and Kaosio Saephan for coordinating the shipment and receipt of samples and Chips Hoai for guiding us regarding regulations related to the import and export of samples. We thank Diana Dresbach for technical help with the trial at the Max Planck Institute. We gratefully acknowledge the financial support from the U.S. Department of Energy (DOE) Office of Science, Office of Biological and Environmental Research for support of the Trial Ecosystem Advancement for Microbiome Science (TEAMS) led by LBNL, Award DE-SC0021234 led by UC San Diego, and m-CAFEs Microbial Community Analysis & Functional Evaluation in Soils, a Science Focus Area led by Lawrence Berkeley National Laboratory where all of the LBNL led projects are under Contract No. DE-AC02-05CH11231. Data analysis utilized resources from the National Energy Research Scientific Computing Center, a DOE Office of Science User Facility (Contract No. DE-AC02-05CH11231). A portion of these data was produced by the US Department of Energy Joint Genome Institute (https://ror.org/04xm1d337; operated under Contract No. DE-AC02-05CH11231). J.C. and M.W. acknowledge support from the University of Melbourne Botany Foundation.

## 8. Data availability

All of the links to the protocols and data as well as the study metadata are available via NMDC at https://data.microbiomedata.org/details/study/nmdc:sty-11-ev70y104. The 16S rRNA amplicon sequencing data are available via NCBI (https://www.ncbi.nlm.nih.gov/) as BioProject PRJNA1151037. All raw data, including plant phenotypes, sterility tests, metabolite identifications, and *in vitro* assays are available via Figshare at https://doi.org/10.6084/m9.figshare.c.7373842. The untargeted metabolomics outputs (HILIC-pos) with features annotations and .mzml files are available via GNPS2 at https://gnps2.org/status?task=2ccbf82840724c99a2acc2c9e512a302. Raw LC-MS/MS files are available in .raw format at MassIVE (https://massive.ucsd.edu/) under ID number MSV000095476 or via https://doi.org/10.25345/C5Q23RB6B. The protocol is available at protocols.io via https://dx.doi.org/10.17504/protocols.io.kxygxyydkl8j/v1. The annotated bacterial genomes can be accessed via IMG/M (https://img.jgi.doe.gov/) by searching for either the isolate name, taxon ID, GOLD Project ID, or by using the links in Table S2.

## 9. Supplementary material

**Table S1:** Chamber settings and models and experiment timing

**Table S2:** Syncom members overview and their OD600 to CFU conversion ratios

**Table S3:** LC-MS parameters

**Table S4:** Metabolite identification and intensity

**Table S5:** Comparison of KEGG modules between SynCom16 vs. Paraburkholderia sp. OAS925

**Video S1:** Z-Stack: RFP-Paraburkholderia motility and colonization on B. distachyon roots at 1 DAI in EcoFAB 2.0

**Video S2:** Root colonization by RFP-Paraburkholderia at 3 DAI in EcoFAB 2.0. Merged bright-field and TxRed channels followed by TxRed footage.

**File S1:** MZMine 3 settings

---

**Fig. S1:**
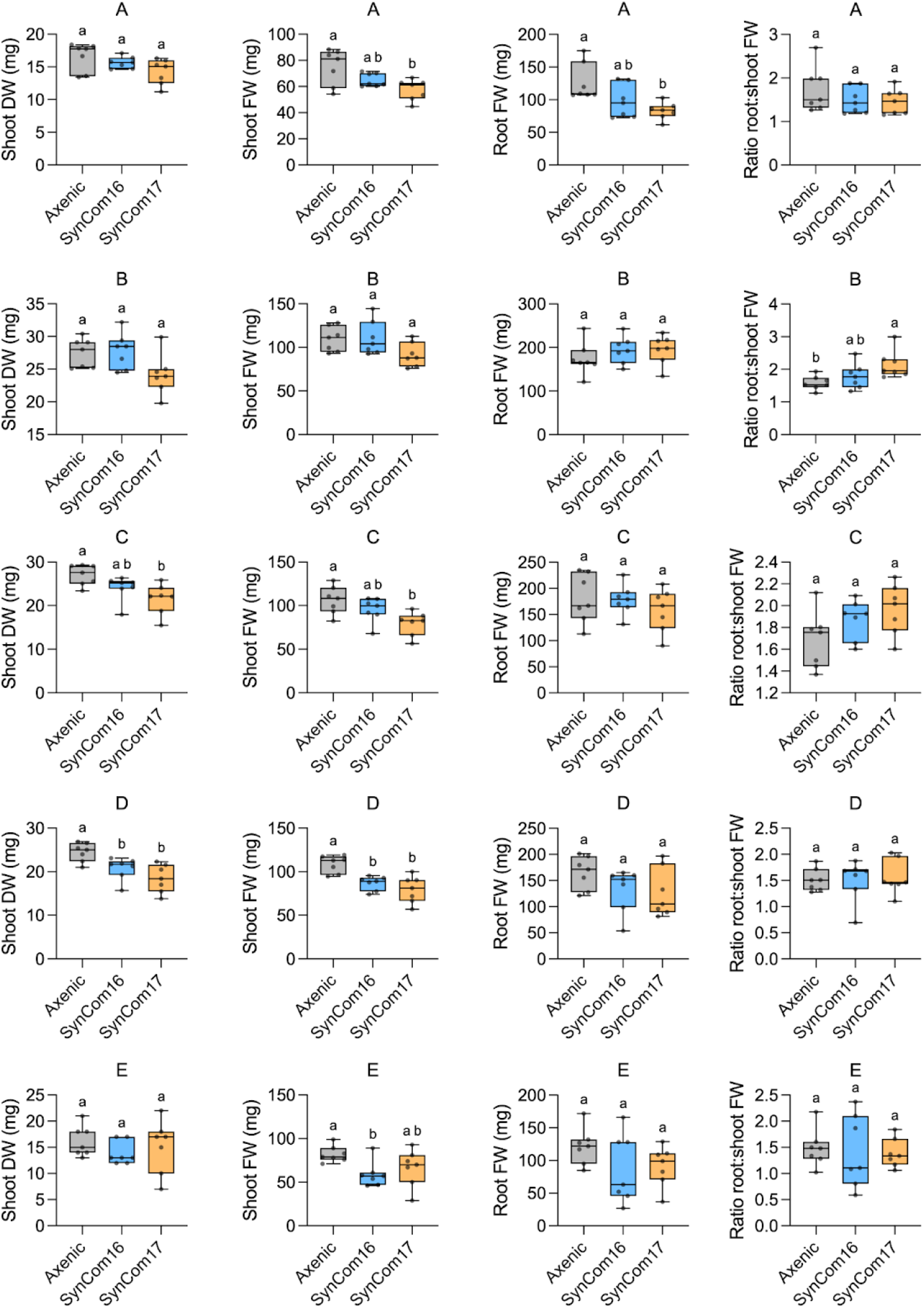
**Plant biomass data for laboratories A-E**. Box plots display all data points, with hinges spanning the 25th to 75th percentiles, a central line denoting the median, and whiskers reaching the minimum and maximum values. Different lowercase letters indicate statistically significant differences at *p*<0.05. One-way ANOVA with Tukey test (*n* = 7).

**Fig. S2:**
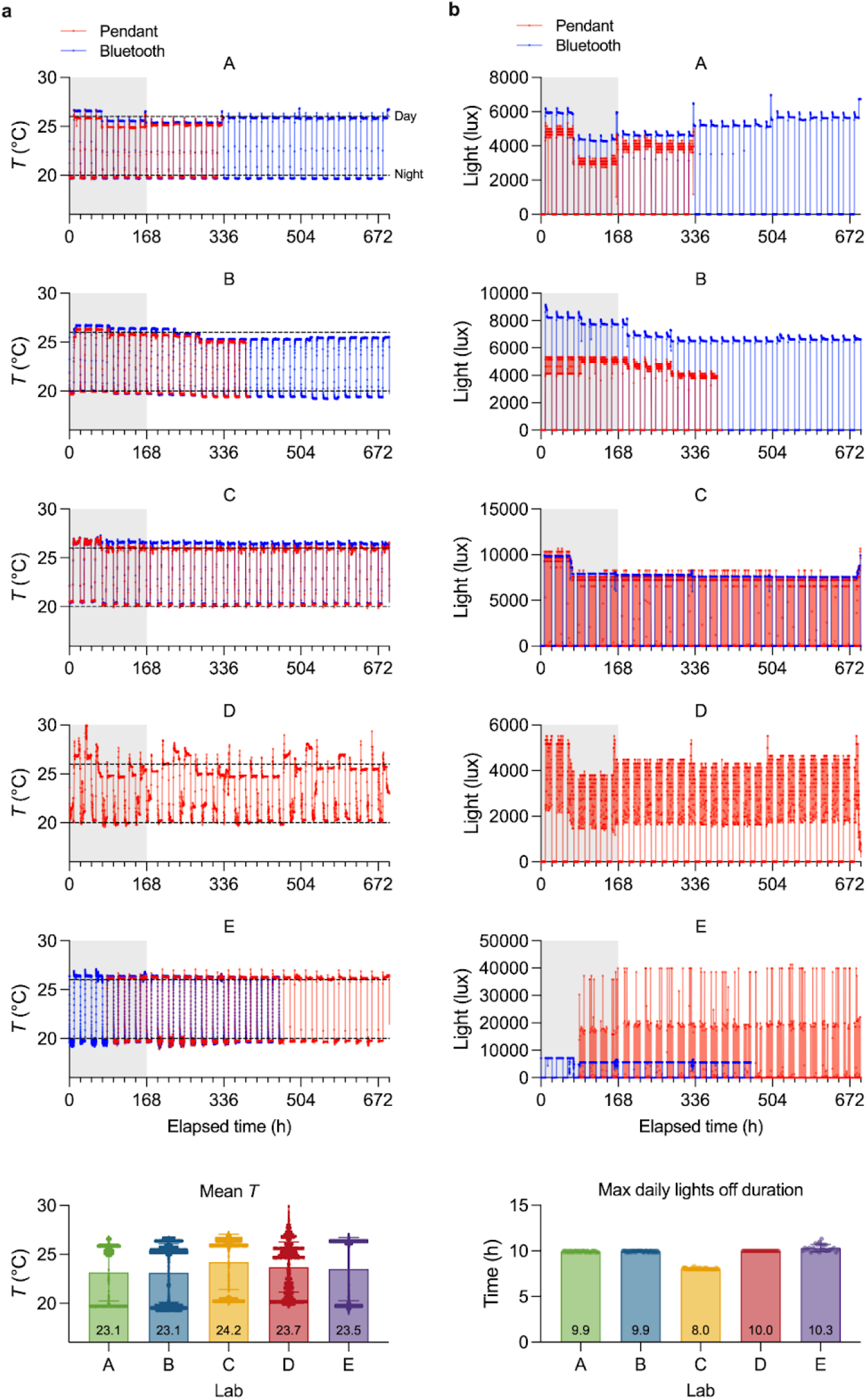
Plant growth conditions in labs A–E: (**a**) temperature (*T*) and (**b**) illuminance to assess lights-off duration, measured by HOBO loggers (Pendant model #UA-00264 in red, Bluetooth model #MX2202 in blue). Dashed lines show the set day/night *T* (26/20°C); gray areas mark the 7-day pre-inoculation period. Labs A, B, and E experienced logging interruptions due to battery drainage; Lab D’s Bluetooth logger did not cover the experimental period.

**Fig. S3:**
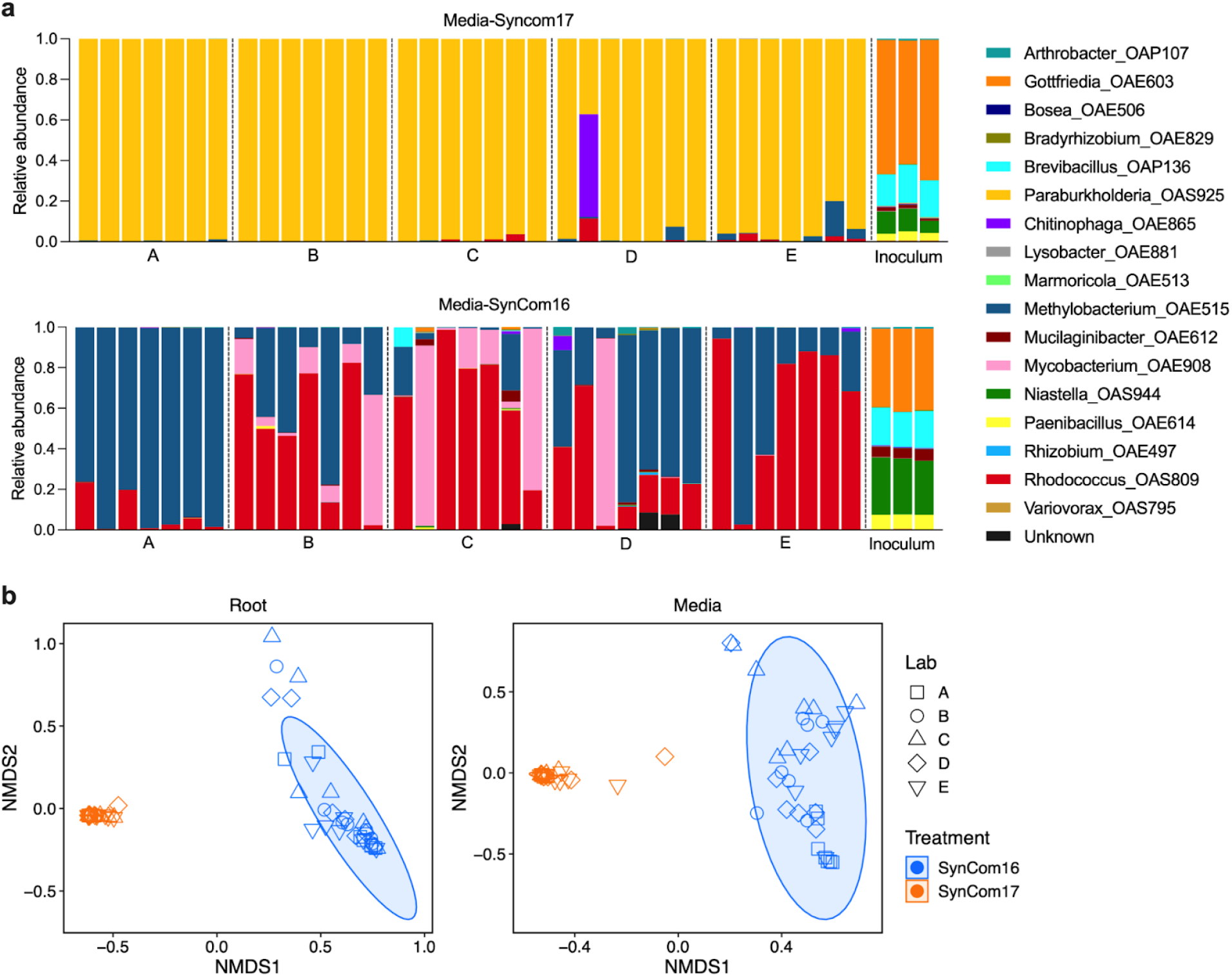
(**a**) **Microbiome composition in plant growth media and starting inoculum**. Letters indicate different laboratories, with each biological replicate shown (*n*=7). The inoculum shows technical replicates (*n*=3). (**b**) NMDS - plot with 95% confidence ellipse. Different laboratories are shown with various symbols, while colors represent SynCom16 (blue) vs. SynCom17 (orange) inoculated plants.

**Fig. S4:**
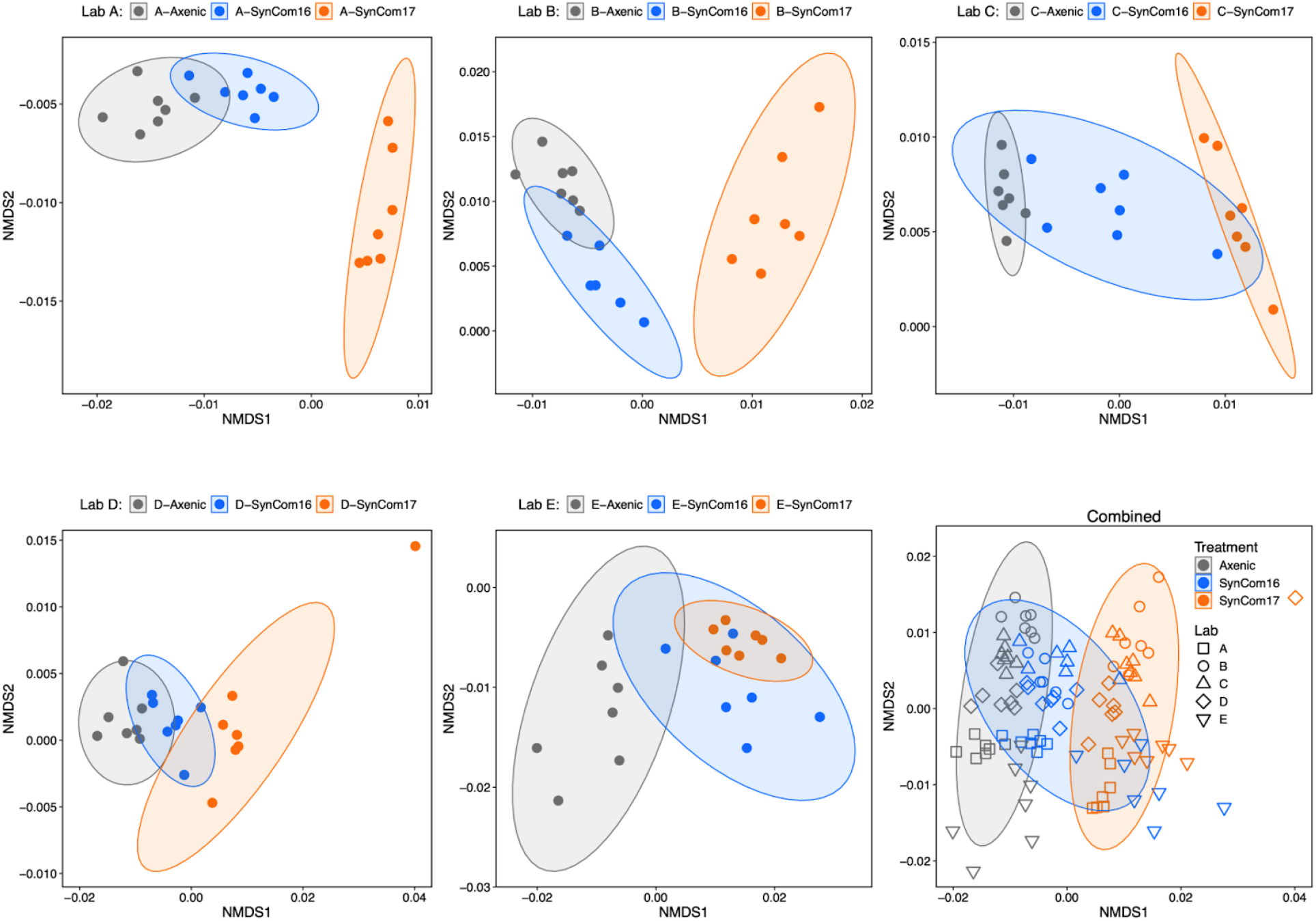
Untargeted metabolomics on root exudates. NMDS plots with a 95% confidence ellipse for 833 filter features for individual laboratories A-E and all combined. Different colors show treatments: Axenic (gray), SynCom16 (blue), and SynCom17 (orange), while shapes indicate laboratories in the combined plot.

**Fig. S5:**
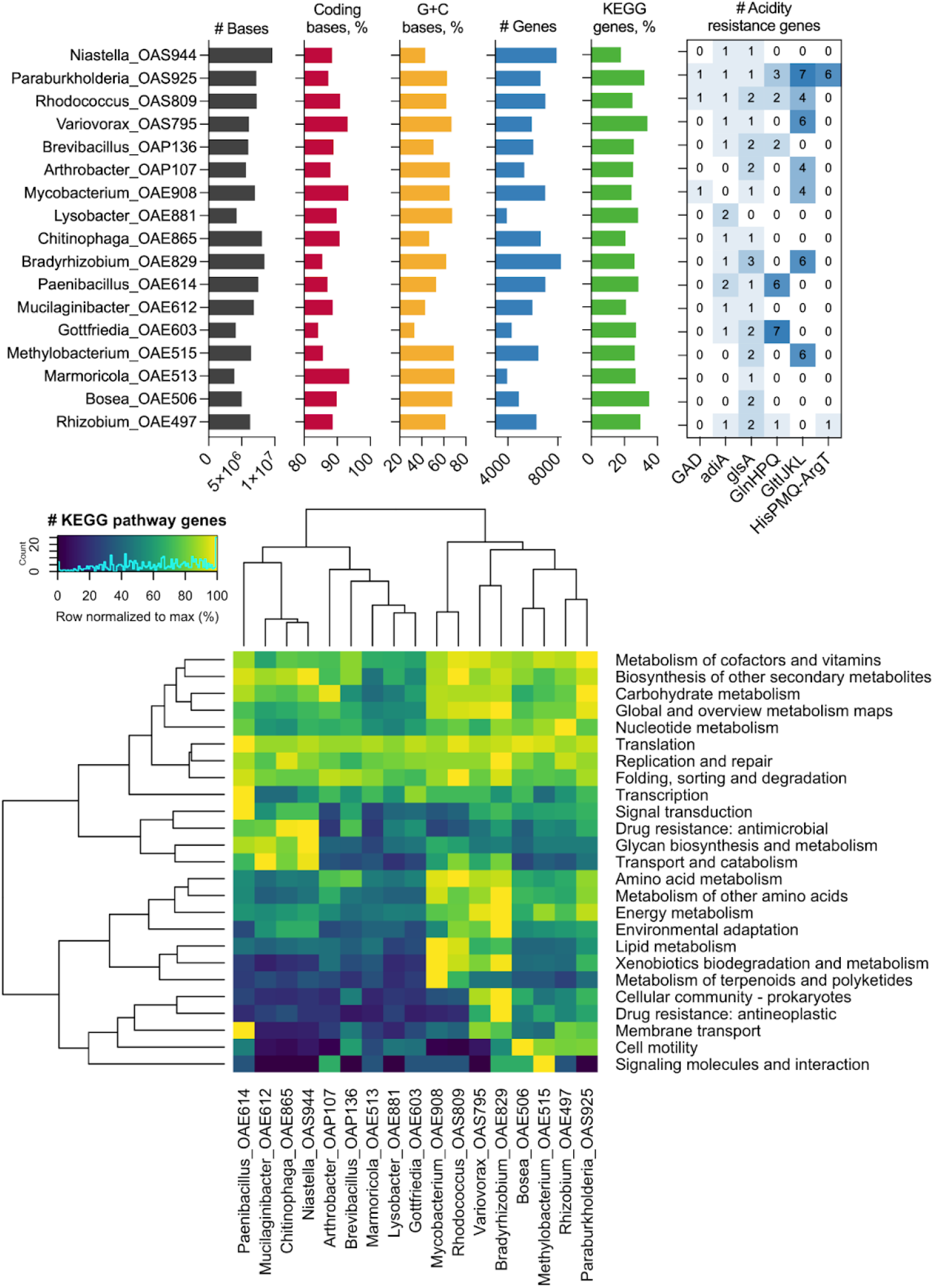
**Comparative genomics of bacterial isolates**. Bar graphs show genome characteristics (from left: the abundance of bases, coding, G+C bases, total genes, and KEGG pathway genes. The search for acid resistance system genes included GAD (EC 4.1.1.15 glutamate decarboxylase), AdiA (EC 4.1.1.19 arginine decarboxylase), glsA (EC 3.5.1.2 glutaminase), GlnHPQ (glutamine ABC transporter), GltIJKL (glutamate/aspartate ABC transporter), HisPMQ-ArgT (arginine/ornithine ABC transporter). The heat map shows normalized gene abundance for selected KEGG pathways.

**Fig. S6:**
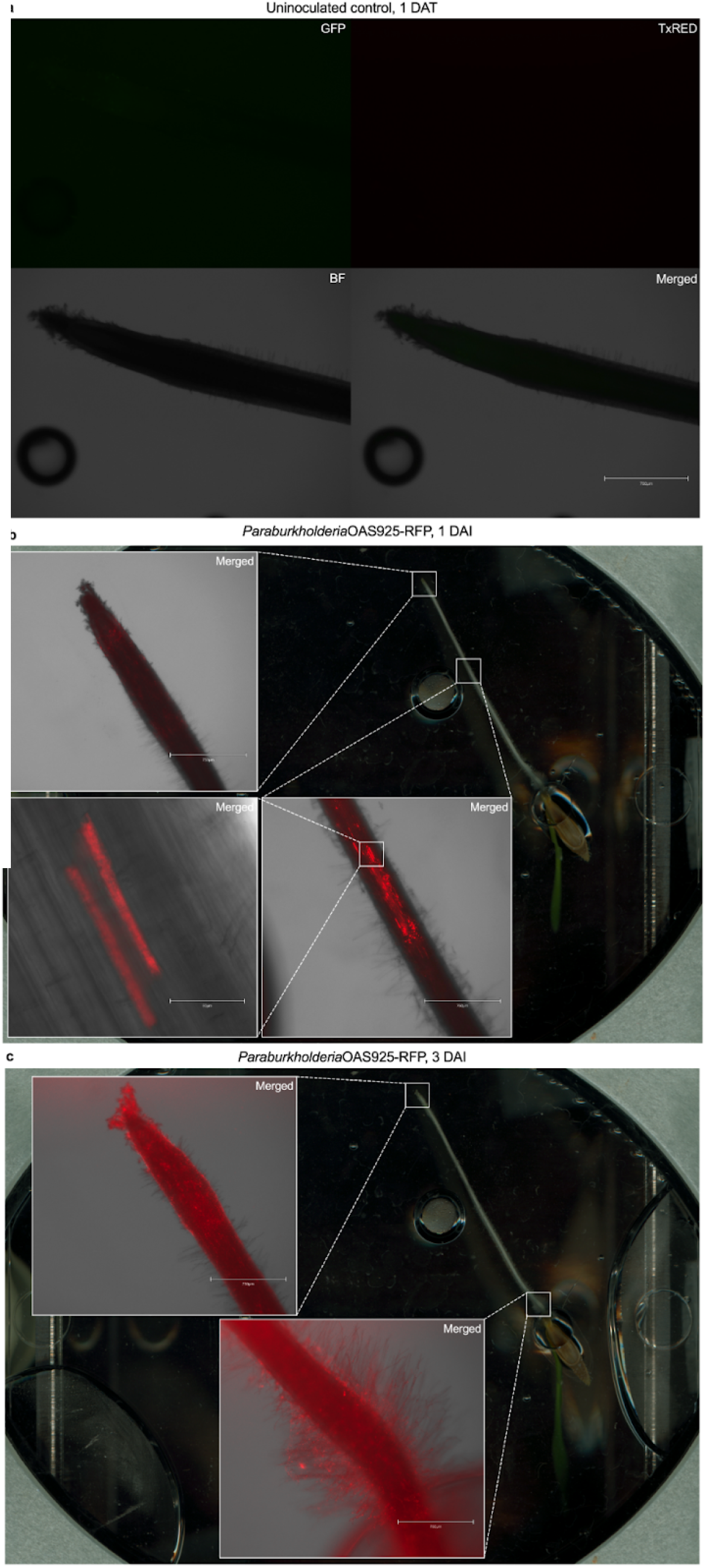
Fluorescent Microscopy in EcoFAB 2.0. We inoculated RFP-expressing *Paraburkholderia* sp. OAS925 into the *B. distachyon* rhizosphere in EcoFAB 2.0 devices on the day of transfer (DAT) for the seedling. The plots show (**a**) uninoculated plant control (1 DAT) and medium-inoculated plants (OD_600_ 0.01) at (**b**) 1 and (**c**) 3 days after inoculation (DAI). EcoFAB 2.0 root scans indicate the locations for microscopy. The inset microscopy images show merged TxRed and bright-field (BF) channels.

**Fig. S7:**
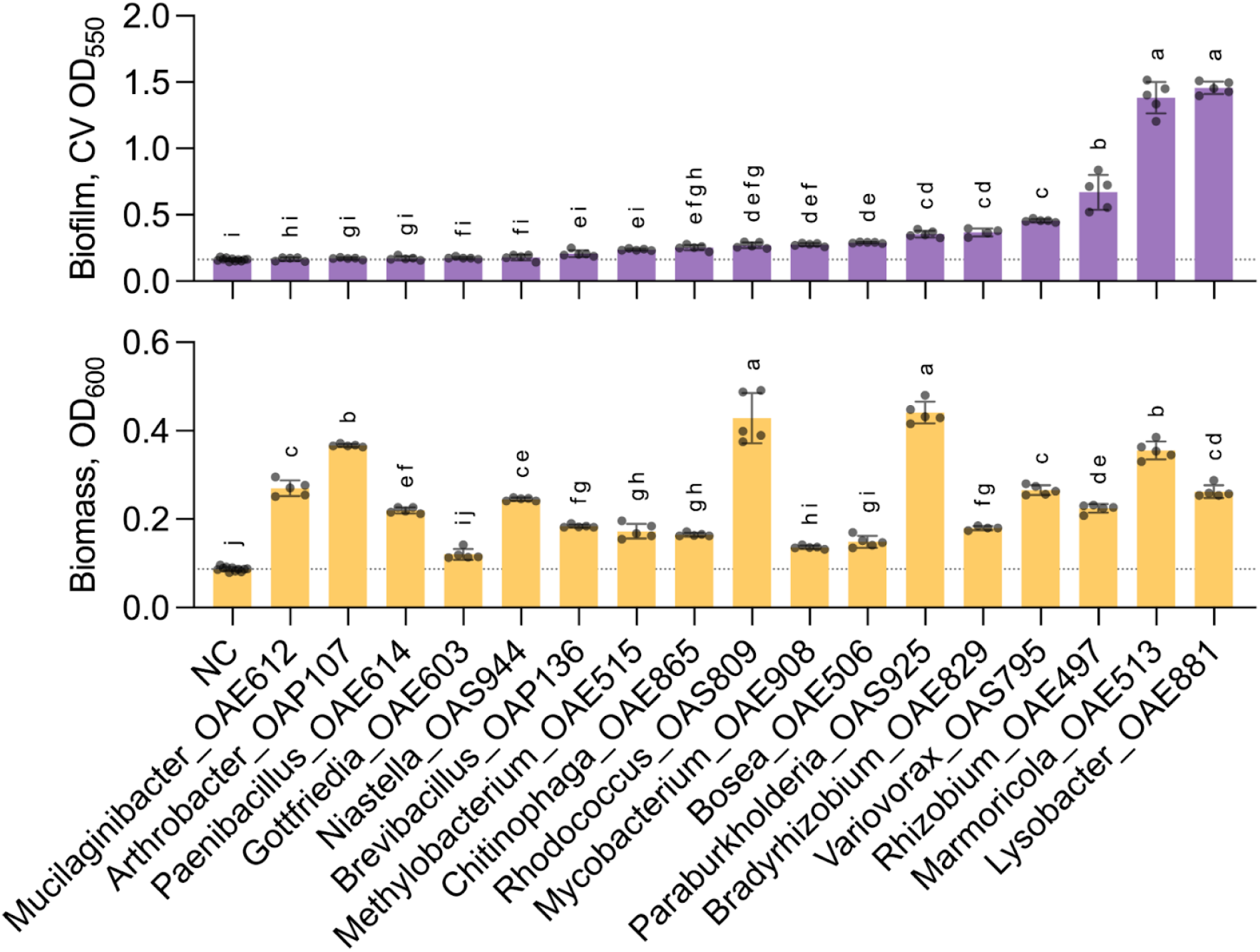
**Biofilm formation for bacterial isolates *in vitro***. Biofilm was measured with crystal violet (CV) staining at OD_550,_ and bacterial biomass was estimated by OD_600_ values of isolates grown on the defined NLDM liquid medium. The horizontal dotted line indicates the mean value of sterile medium negative control (NC). Different letters indicate statistically significant differences at *p*<0.05, One-way ANOVA with Tukey’s test, *n*=4–5 for isolates and *n*=11 for NC.

**Fig. S8:**
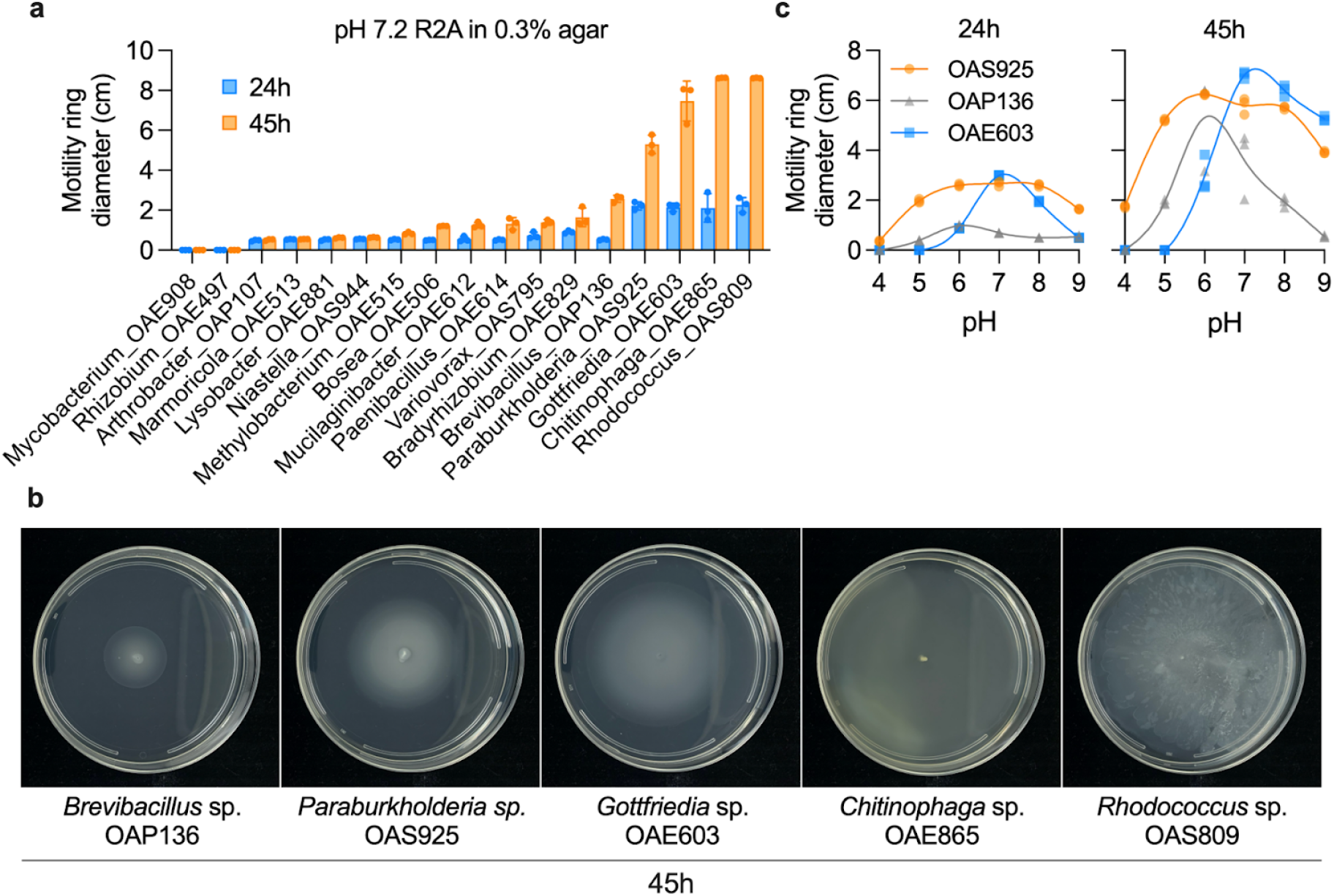
**Motility assays**. (**a**) The initial screen for swimming motility across bacterial isolates was measured 24 and 45 h after inoculation. (**b**) Phenotypes of the most motile strains at 45 h since inoculation. (**c**) pH effects on the motility ring diameter of isolates with bulls-eye colony morphology.

## References

1. Zengler, K. et al. EcoFABs: advancing microbiome science through standardized fabricated ecosystems. Nat. Methods 16, 567–571 (2019).

2. O’Toole, G. A. We have a community problem. J. Bacteriol. 206, e0007324 (2024).

3. Vorholt, J. A., Vogel, C., Carlström, C. I. & Müller, D. B. Establishing causality: opportunities of synthetic communities for plant microbiome research. Cell Host Microbe 22, 142–155 (2017).

4. Marín, O., González, B. & Poupin, M. J. From microbial dynamics to functionality in the rhizosphere: A systematic review of the opportunities with synthetic microbial communities. Front. Plant Sci. 12, 650609 (2021).

5. Salas-González, I. et al. Coordination between microbiota and root endodermis supports plant mineral nutrient homeostasis. Science 371, (2021).

6. Finkel, O. M. et al. A single bacterial genus maintains root growth in a complex microbiome. Nature 587, 103–108 (2020).

7. Vandenkoornhuyse, P., Quaiser, A., Duhamel, M., Le Van, A. & Dufresne, A. The importance of the microbiome of the plant holobiont. New Phytol. 206, 1196–1206 (2015).

8. Northen, T. R., et al. Community standards and future opportunities for synthetic communities in plant-microbiota research (in press). Nat Microbiol (2024).

9. Coker, J., et al. A Reproducible and Tunable Synthetic Soil Microbial Community Provides New Insights into Microbial Ecology. mSystems 7, e0095122 (2022).

10. Sasse, J. et al. Multilab EcoFAB study shows highly reproducible physiology and depletion of soil metabolites by a model grass. New Phytol. 222, 1149–1160 (2019).

11. Lin, H.-H. et al. Impact of inoculation practices on microbiota assembly and community stability in a fabricated ecosystem. Phytobiomes Journal (2023) doi:10.1094/PBIOMES-06-23-0050-R.

12. Novak, V. et al. Reproducible growth of Brachypodium in EcoFAB 2.0 reveals that nitrogen form and starvation modulate root exudation. Sci. Adv. 10, eadg7888 (2024).

13. McGrath, T. F., Haughey, S. A., Islam, M., Elliott, C. T. & Collaborators: The potential of handheld near infrared spectroscopy to detect food adulteration: Results of a global, multi-instrument inter-laboratory study. Food Chem. 353, 128718 (2021).

14. Thompson, M., Ellison, S. L. R. & Wood, R. The International Harmonized Protocol for the proficiency testing of analytical chemistry laboratories (IUPAC Technical Report). Pure Appl. Chem. 78, 145–196 (2006).

15. Zhalnina, K. et al. Dynamic root exudate chemistry and microbial substrate preferences drive patterns in rhizosphere microbial community assembly. Nat. Microbiol. 3, 470–480 (2018).

16. Korenblum, E. et al. Rhizosphere microbiome mediates systemic root metabolite exudation by root-to-root signaling. Proc Natl Acad Sci USA 117, 3874–3883 (2020).

17. Allison, S. D. & Vitousek, P. M. Responses of extracellular enzymes to simple and complex nutrient inputs. Soil Biol. Biochem 37, 937–944 (2005).

18. de Raad, M. et al. A defined medium for cultivation and exometabolite profiling of soil bacteria. Front. Microbiol. 13, 855331 (2022).

19. Schloss, P. D. Identifying and overcoming threats to reproducibility, replicability, robustness, and generalizability in microbiome research. MBio 9, (2018).

20. Nuccio, E. E. et al. Niche differentiation is spatially and temporally regulated in the rhizosphere. ISME J. 14, 999–1014 (2020).

21. Fierer, N. Embracing the unknown: disentangling the complexities of the soil microbiome. Nat. Rev. Microbiol. 15, 579–590 (2017).

22. Debray, R. et al. Priority effects in microbiome assembly. Nat. Rev. Microbiol. 20, 109–121 (2022).

23. Carlström, C. I. et al. Synthetic microbiota reveal priority effects and keystone strains in the Arabidopsis phyllosphere. *Nat*. Ecol. Evol. 3, 1445–1454 (2019).

24. Kuzyakov, Y. & Razavi, B. S. Rhizosphere size and shape: Temporal dynamics and spatial stationarity. Soil Biol. Biochem 135, 343–360 (2019).

25. Lu, P. et al. L-glutamine provides acid resistance for Escherichia coli through enzymatic release of ammonia. Cell Res. 23, 635–644 (2013).

26. Minamino, T., Imae, Y., Oosawa, F., Kobayashi, Y. & Oosawa, K. Effect of intracellular pH on rotational speed of bacterial flagellar motors. J. Bacteriol. 185, 1190–1194 (2003).

27. Cole, B. J. et al. Genome-wide identification of bacterial plant colonization genes. PLoS Biol. 15, e2002860 (2017).

28. Hafner, K., Welt, B. & Pelletier, W. Dry ice sublimation performance as affected by binding agent, density, and age. Engineering … (2023).

29. Teytelman, L., Stoliartchouk, A., Kindler, L. & Hurwitz, B. L. Protocols.io: virtual communities for protocol development and discussion. PLoS Biol. 14, e1002538 (2016).

30. Eloe-Fadrosh, E. A. et al. The National Microbiome Data Collaborative Data Portal: an integrated multi-omics microbiome data resource. NAT (2022) doi:10.1093/nar/gkab990.

31. Sinha, R. et al. Assessment of variation in microbial community amplicon sequencing by the Microbiome Quality Control (MBQC) project consortium. Nat. Biotechnol. 35, 1077–1086 (2017).

32. Vogel, J. & Hill, T. High-efficiency *Agrobacterium*-mediated transformation of *Brachypodium distachyon* inbred line Bd21-3. Plant Cell Rep. 27, 471–478 (2008).

33. Zavafer, A., Mancilla, C., Jolley, G. & Murakami, K. On the concepts and correct use of radiometric quantities for assessing the light environment and their application to plant research. Biophys. Rev. 15, 385–400 (2023).

34. Sordo, Z., Andeer, P., Sethian, J., Northen, T. & Ushizima, D. RhizoNet segments plant roots to assess biomass and growth for enabling self-driving labs. Sci. Rep. 14, 12907 (2024).

35. Schindelin, J. et al. Fiji: an open-source platform for biological-image analysis. Nat. Methods 9, 676–682 (2012).

36. Lobet, G., Pagès, L. & Draye, X. A novel image-analysis toolbox enabling quantitative analysis of root system architecture. Plant Physiol. 157, 29–39 (2011).

37. Hulstaert, N. et al. ThermoRawFileParser: Modular, Scalable, and Cross-Platform RAW File Conversion. J. Proteome Res. 19, 537–542 (2020).

38. Schmid, R. et al. Integrative analysis of multimodal mass spectrometry data in MZmine 3. Nat. Biotechnol. 41, 447–449 (2023).

39. Wang, M. et al. Sharing and community curation of mass spectrometry data with Global Natural Products Social Molecular Networking. Nat. Biotechnol. 34, 828–837 (2016).

40. Sumner, L. W. et al. Proposed minimum reporting standards for chemical analysis Chemical Analysis Working Group (CAWG) Metabolomics Standards Initiative (MSI). Metabolomics 3, 211–221 (2007).

41. Bowen, B. P. & Northen, T. R. Dealing with the unknown: metabolomics and metabolite atlases. J. Am. Soc. Mass Spectrom. 21, 1471–1476 (2010).

42. Kim, S. Exploring chemical information in pubchem. Curr. Protoc. 1, e217 (2021).

43. Parada, A. E., Needham, D. M. & Fuhrman, J. A. Every base matters: assessing small subunit rRNA primers for marine microbiomes with mock communities, time series and global field samples. Environ. Microbiol. 18, 1403–1414 (2016).

44. Apprill, A., McNally, S., Parsons, R. & Weber, L. Minor revision to V4 region SSU rRNA 806R gene primer greatly increases detection of SAR11 bacterioplankton. Aquat. Microb. Ecol. 75, 129–137 (2015).

45. Edgar, R. C. Search and clustering orders of magnitude faster than BLAST. Bioinformatics 26, 2460–2461 (2010).

46. Haney, E. F., Trimble, M. J. & Hancock, R. E. W. Microtiter plate assays to assess antibiofilm activity against bacteria. Nat. Protoc. 16, 2615–2632 (2021).

47. Mukherjee, S. et al. Twenty-five years of Genomes OnLine Database (GOLD): data updates and new features in v.9. Nucleic Acids Res. 51, D957–D963 (2023).

48. Chen, I.-M. A. et al. The IMG/M data management and analysis system v.7: content updates and new features. Nucleic Acids Res. 51, D723–D732 (2023).

49. Chen, I.-M. A. et al. The IMG/M data management and analysis system v.6.0: new tools and advanced capabilities. Nucleic Acids Res. 49, D751–D763 (2021).

50. Parks, D. H. et al. GTDB: an ongoing census of bacterial and archaeal diversity through a phylogenetically consistent, rank normalized and complete genome-based taxonomy. Nucleic Acids Res. 50, D785–D794 (2022).

51. Price, M. N., Dehal, P. S. & Arkin, A. P. FastTree 2 — approximately maximum-likelihood trees for large alignments. PLoS ONE 5, e9490 (2010).

52. Letunic, I. & Bork, P. Interactive Tree Of Life (iTOL) v5: an online tool for phylogenetic tree display and annotation. Nucleic Acids Res. 49, W293–W296 (2021).

53. Sayers, E. W. et al. Database resources of the National Center for Biotechnology Information. Nucleic Acids Res. 39, D38–51 (2011).

54. Pearson, A. N. et al. The pGinger Family of Expression Plasmids. Microbiol. Spectr. 11, e0037323 (2023).

55. Kearns, D. B. A field guide to bacterial swarming motility. Nat. Rev. Microbiol. 8, 634–644 (2010).

56. Warnes, G. R., et al. gplots: Various R Programming Tools for Plotting Data. R package version 3.1.3.1. https://CRAN.R-project.org/package=gplots (2024).

